# *Basella alba* L. var. ‘Rubra’ L-DOPA/dopamine-4,5-dioxygenase 1 prefers L-DOPA over dopamine and ascorbic acid enhances its activity

**DOI:** 10.64898/2026.02.19.706318

**Authors:** Hidam Bishworjit Singh, Mohammad Imtiyaj Khan

**Affiliations:** Biochemistry and Molecular Biology Lab, Department of Biotechnology, Gauhati University, Assam, India

**Keywords:** RACE-PCR, enzyme activation, L-DOPA, dopamine, ascorbic acid, crowding effect, phylogeny

## Abstract

Betalamic acid is the chromophore of betalains, which are pigments of chemotaxonomic and physiological importance. This study involves RACE-PCR-based gene cloning, heterologous expression, protein purification, and steady-state kinetics of *B. alba* L. var. ‘Rubra’ L-DOPA/dopamine-4,5-dioxygenase 1 (BrDOD1). BrDOD1 is a unique high betalamic acid-forming LigB homolog in plants having comparable affinity for both L-DOPA and dopamine (*K*_M_ < 50 *μ*M). Ascorbic acid (10 mM) shifted the steady-state kinetics from inhibitory to activator at a particular substrate concentration of both L-DOPA and dopamine. This increased both *K*_M_ and *V*_max_ by more than 6.5-fold, indicating that ascorbic acid acted as a molecular crowding agent in the enzyme assay. BrDOD1’s physiological substrate is L-DOPA as the reaction rate for L-DOPA was 6.6-fold higher than dopamine, L-DOPA was present in higher concentration than that of dopamine in the same plant, and molecular dynamic simulations showed better stability of BrDOD1−L-DOPA complex than that of dopamine. Further, two more LigB homologs from the same plant have also been cloned. Based on the betalamic acid-forming activity, molecular phylogeny, conserved structural regions, and theoretical pI, betalainic plant LigB homologs have been classified into three groups to better understand the evolutionary trajectory of the LigB homologs in plants.

## 1. Introduction

L-3,4-dihydroxyphenylalanine (L-DOPA) dioxygenases are broadly of two types: L-DOPA- 2,3-and 4,5-specific dioxygenase (or DODA, which is present in some microbes and fungi) and L-DOPA-4,5-dioxygenase (or DOD reported from betalainic plants). DODAs and DODs are extradiol-type aromatic ring-opening enzymes, which are characterized by two His residues and one Glu/Asp residue involved in the catalysis (Burroughs et al., 2019; Christinet et al., 2004; Kovaleva and Lipscomb, 2007). Such aromatic ring-opening reactions involving oxidation by dioxygenases are generally parts of degradation pathways, particularly that of lignin-derived aromatic compounds (LDAC). Both DODAs and DODs are homologs of LigB, which is distributed across different kingdoms of living beings. LigB is part of the enzyme system involved in the breakdown of lignin in bacteria (Noda et al., 1990; Spence et al., 1996; Sugimoto et al., 1999). The aromatic ring-opening reactions of DODA and DOD resemble that of the degradation of LDAC by LigAB in bacteria (Barry and Taylor, 2013). The LigB homolog present in *Portulaca grandiflora* (PgDOD) contains the bacterial LigB domain (Christinet et al., 2004). Interestingly, these LigB homologs, DODAs and DODs, are responsible for the formation of health-promoting water-soluble plant pigments – betalains – that are used for coloring foods, cosmetics, smart indicators, and fluorescent probes (Khan, 2016; Khan and Liu, 2022; Khan and Polturak, 2025; Martínez-Rodríguez et al., 2024). These pigments have important functional roles *in planta* (Davies et al., 2022), possibly associated with pleiotropic effect on certain phenotype (Kumari and Khan, 2025), and other organisms in protection against radiation (Li et al., 2024). In addition, these pigments are also chemotaxonomic markers in a number of plants of the Caryophyllales order because of their mutual exclusiveness with anthocyanins(Khan, 2016; Pucker et al., 2024; Waterman, 2007). In betalain biosynthesis of plants, the rate-limiting step is catalyzed by DOD (Jung and Maeda, 2024) through the aromatic ring-opening reaction of L-DOPA cleaving at C-4,5 to ultimately produce betalamic acid (BA) (Fig. S1). The same reaction is catalyzed by DODA in prokaryotes and fungi (Contreras-Llano et al., 2019; Mueller et al., 1997a). However, unlike DOD, DODA can produce BA and muscaflavin through the aromatic ring-opening of L-DOPA by cleaving L-DOPA at C-4,5 and C-2,3, respectively. The first DODA coding cDNA was cloned from *Amanita muscaria* (*AmDODA*) and shown through heterologous expression to be bifunctional (Hinz et al., 1997). Following this, many homologs have been cloned from betalain-accumulating plants and functionally validated through heterologous expression (Chou et al., 2019; Christinet et al., 2004; Chung et al., 2015; Gandía-Herrero and García-Carmona, 2012; Henarejos-Escudero et al., 2022; Sasaki et al., 2009; Sheehan et al., 2020). In addition, in the above-mentioned studies, from the same betalain-accumulating plant, multiple DOD paralogs were reported, with some showing involvement in betalain biosynthesis but with varying activities, i.e., high or low activity and differential expression, while others showing no such functional role, i.e., they do not cleave the aromatic ring in L-DOPA to form BA (Brockington et al., 2015; Sheehan et al., 2020). *LigB* homologs have been reported in non-betalainic plants also, but they do not exhibit BA-forming activity (Brockington et al., 2015; Kasei et al., 2021; Li et al., 2025; Sheehan et al., 2020; Weng et al., 2012). However, the *LigB* gene from non-betalainic plant *Arabidopsis thaliana* has been implicated in the formation of arabidopyrones derived from caffealdehyde (Weng et al., 2012) or caffeic acid (Kasei et al., 2021). Similarly, pansy (*Viola* × *wittrockiana*) LigB homolog also has been shown to produce arabidopyrones from caffeic acid to confer stress tolerance by improving the antioxidant system (Li et al., 2025).

Emerging evidence points toward possible dual-activity of plant DODs by acting on both L-DOPA and dopamine, which are structurally related. *Mirabilis jalapa* DOD (MjDOD1) was used successfully in a whole-cell biosensor to detect dopamine, indicating that the enzyme can act on both L-DOPA and dopamine (Lin and Yeh, 2017). The dual substrate affinity was also observed in *Chenopodium quinoa* DOD, with a higher affinity for dopamine than L-DOPA (Henarejos-Escudero et al., 2022). As kinetic parameters of only three high BA- forming activity plant DODs have been reported so far (Chou et al., 2019; Guerrero-Rubio et al., 2023; Henarejos-Escudero et al., 2022), there is scant information on the dual substrate affinity of other plant DODs. Since DOD catalyzes the rate-limiting step in the biosynthetic pathway of betalains and L-DOPA and dopamine are produced from the same precursor, tyrosine, it is of utmost importance to study the kinetic parameters of other plant DODs to understand the substrate preference of the enzyme vis-a-vis the ubiquitous reducing agent, ascorbic acid, which has been used commonly in all the DOD kinetic studies so far. In the present study, we picked a gomphrenin-producing plant, *Basella alba* L. var. ‘Rubra’ to study the kinetics of DOD enzyme. *B. alba* L. var. ‘Rubra’ is a heat-tolerant and fast-growing leafy vegetable abundant in tropical Asia and Africa. It is rich in minerals, vitamins, and antioxidants, effectively treating various diseases and offering health benefits (Chaurasiya et al., 2021). Also, it is a significant source of betalains, with gomphrenin I being the predominant pigment (Lin et al., 2010). While numerous studies have reported on the phytochemicals and pigment analysis of *Basella* spp., the molecular analysis and biochemical characterization of the genes involved in the biosynthesis of its major pigment, betalains, are yet to be conducted. Herein, we report the cloning and characterization of steady-state kinetics of *B. alba* L. var. ‘Rubra’ DOD1 (BrDOD1) using L-DOPA and dopamine as substrates. Further, we also report for the first time the differential effects of ascorbic acid on DOD kinetics against L-DOPA and dopamine, and identify the possible physiological substrate of BrDOD1. Furthermore, two more *LigB* homologs from *B. alba* L. var. ‘Rubra’ have been mined, and all the three LigB homologs have been classified based on BA-forming activity and other characteristic features.

## 2. Materials and Method

### 2.1. Strains and reagents

*E. coli* strains DH5α and BL21-CodonPlus were provided by Dr. P. Giridhar of CSIR-CFTRI, Mysuru, India and Dr Ashok K. Varma of ACTREC, Mumbai, India, respectively. All the analytical-grade reagents, chemicals, HPLC-grade solvents, molecular biology enzymes, and kits were purchased from Merck (Darmstadt, Germany), Sigma-Aldrich (Massachusetts, USA), Sisco Research Laboratory (Mumbai, India), Thermo Fischer Scientific (Massachusetts, USA), or HiMedia (Mumbai, India).

### 2.2. Gene mining and cloning

To amplify the partial gene sequences of LigB homologs, primers were designed (Table S1A) using the IDT OligoAnalyzer tool (https://www.idtdna.com/pages/tools/ OligoAnalyzer). Total RNA from *B. alba* L. var. ‘Rubra’ leaf (Fig. S2) was isolated using the CTAB-LiCl_2_ method, and then cDNA was prepared from 2 *µ*g of RNA using RevertAid cDNA synthesis kit (Thermo Scientific). Then, the partial sequences of LigB homologs were amplified in a thermal cycler (Applied Biosystem, Massachusetts, USA) under optimized conditions. The gene fragments were then purified using the GeneJET gel extraction kit (Thermo Scientific), followed by ligation to the pJET1.2 blunt vector (Thermo Scientific) and transformation into *E. coli* DH5α using the CaCl_2_ heat-shock method. The transformed cells were allowed to grow on ampicillin-LB agar plates at 37 °C overnight. Transformed colonies were selected and allowed to grow in LB broth, Miller (HiMedia), supplemented with 100 *µ*g/mL ampicillin (SRL), then incubated in a shaking incubator at 37 °C overnight. Plasmids were then isolated using alkaline lysis method and confirmed through double restriction digestion and Sanger sequencing. The partial gene sequence of the LigB homolog was validated by searching the NCBI database using the NCBI-BLAST tool. For the full-length gene amplification, nested PCR primers (Table S1B) specific to the validated partial gene sequence were designed using the IDT-OligoAnalyzer tool to amplify both the unknown regions at the 3′ and 5′ ends flanking the known partial sequence using rapid amplification of cDNA ends PCR (RACE-PCR). Then, PCR was performed to amplify the 3′ and 5′ ends of the partial gene sequence from cDNA using the RACE PCR kit (Sigma) per the manufacturer’s protocol under the optimized PCR conditions. Amplified 3′ and 5′ end fragments were subsequently cloned and verified as described above. Following confirmation, the sequences were aligned using ClustalX2, and the output alignment file was assembled using the contig assembly program, CAP3, on Bioedit 7.7.1. Using the assembled sequence, the putative open reading frame (ORF) was identified using NCBI homology search and ORF finder tool. Following this, forward and reverse primers (Table S1C) for the full-length amplification of the ORF of DOD1-like gene were designed using the IDT OligoAnalyzer tool. Then, the full-length DOD1-like gene sequence was amplified from the *B. alba* L. var. ‘Rubra’ cDNA under optimized PCR conditions. The product was purified, ligated to pLATE51 expression vector according to the manufacturer’s protocol (LIC cloning kit, Thermo Scientific), and transformed into *E*. *coli* DH5α cells. Transformation confirmation was achieved through plasmid extraction, Sanger sequencing and validation, as described above.

### 2.3. Recombinant protein expression, extraction, and purification

pLATE51::*BrDOD1* expression constructs were mobilized into *E. coli* BL21-CodonPlus cells using the CaCl_2_ heat shock method. Recombinant cells carrying the pLATE51::*BrDOD1* construct were grown in M9 medium supplemented with chloramphenicol and ampicillin. The cells were allowed to grow till the OD reached 0.6 units. Then, the cells were induced by adding the final concentration of 1 mM isopropyl β-D-1-thiogalactopyranoside (IPTG; HiMedia) and allowed to grow overnight in shaking incubator at 37 °C. Overnight culture of 1 mM IPTG-induced *E. coli* BL21-CodonPlus recombinant cells carrying pLATE51::BrDOD1 construct were collected by centrifugation at 5000 ×*g* for 5 min. Then, the cell pellets were washed with phosphate buffered saline (pH 7.2), and the crude protein was extracted by lysing the cells with three cycles of freeze-thaw method in a cell lysis buffer (300 mM NaCl, 50 mM NaH_2_PO_4_ and 10 mM imidazole, adjusted to pH 8.0 with NaOH) supplemented with lysozyme at a final concentration of 1 mg/mL. To reduce the viscosity of the cell lysate caused by nucleic acids, DNase and RNase were added and incubated at ambient temperature for 30 min. The cell lysate was then passed through a 24G syringe, and the supernatant containing crude protein was then collected by centrifugation at 8000 ×*g* for 10 min. The recombinant protein was purified using Ni-NTA affinity chromatography (Qiagen, Hilden, Germany) through a gravity flow column according to the manufacturer’s protocol. Ni-NTA purified protein was stored at 4 °C after flushing with N_2_ gas. The total protein content was estimated using the Bradford method using bovine serum albumin (BSA) standard curve.

### 2.4. Enzyme activation and desalting

The Ni-NTA affinity column purified recombinant enzyme was reconstituted with 10 mM dithiothreitol, 10 mM ascorbic acid, and five times molar concentration of FeSO_4_.7H_2_O and incubated at 4 °C for 30 min after flushing N_2_ gas. After that, all the reconstitution components were removed using a gravity flow PD10 column (M/s Cytiva, Massachusetts, US), and the buffer was exchanged by passing 20 mM phosphate buffer pH 6.5. Desalted and activated enzyme was stored at 4 °C after flushing with N_2_ gas to minimize exposure to atmospheric O_2_. The optimum pH for BrDOD1 was determined in different pH ranges using 50 mM acetate (4.0 – 5.6), phosphate (5.8 – 7.8), and tris-HCl buffer (8.0 - 8.5) at 25 ± 2 °C, and the data for the formation of BA were fitted using a non-linear curve fit in GraphPad Prism 8 (trial subscription).

### 2.5. Spectrophotometric analysis of betalain and reactions product

Betalain pigment quantification (detailed protocol in the supplementary method file section 1.1.) was done using UV-Vis spectrophotometer (Model UV 1800, M/s Shimadzu Corp., Kyoto, Japan) at 535, 480, and 600 nm. Betacyanin, betaxanthin and total betalain content was calculated using the following formula (Khan et al., 2012).

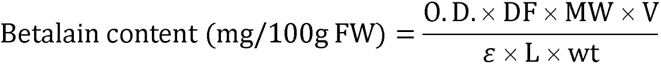

In addition to this, substrate L-DOPA (Sigma) or dopamine (SRL) was added to the purified enzyme assay mixture and incubated for 30 min, and the greenish-yellow color developed in the culture medium was observed and characterized using a spectrophotometer (Shimadzu).

### 2.6. HPLC analysis of pigment and quantification of metabolites

The reaction products were pooled and concentrated by lyophilization. Then, the dried powder was reconstituted with solvent A (0.2% formic acid in water), followed by filtration using a 0.2 *µ*m syringe filter. Aliquots of 20 μL of sample were injected in HPLC (Prominence LC 20AD, Shimadzu Corp., Kyoto, Japan) equipped with a photo-diode array (PDA) detector and separated in a YMC reversed phase column (250×4.6 mm, 5 μm particle size and 12 nm pore size). The reaction mixture was analyzed at a 1.0 mL/min flow rate with a linear gradient of solvent B (acetonitrile) from 7% to 49% over 16 min and then to 7% over 4 min. Furthermore, for metabolite quantification, 20 μL of sample preparations were injected in HPLC (detailed protocol in the supplementary method file section 1.2.). And the quantification was done using standard calibration curves of L-DOPA, dopamine, and ascorbic acid.

### 2.7. Determination of steady-state kinetics and data analysis

To determine the kinetic parameters, a reaction mixture containing 50 mM phosphate buffer pH 6.8, 10 mM ascorbic acid, 2.93 *µ*mole of enzyme, and different concentration ranges of L-DOPA or dopamine (12.5 *µ*M to 3200 *µ*M) were mixed in a total of 350 μL reaction volume and then the rates of formation of BA and 6-decarboxy-BA were recorded in the 96- well multi-scan spectrophotometer (M/s ThermoScientific, Massachusetts, USA) at 430 nm for 30 min. Further, another kinetic analysis was also performed using the same method but without 10 mM ascorbic acid. Thus, the rates of product formation and steady-state kinetics were recorded in the presence and absence of cosolute ascorbic acid in the reaction mixture. The steady-state Michaelis-Menten kinetic data were plotted using a non-linear regression curve using the default best-fit model on GraphPad Prism8 for the reactions containing ascorbic acid. For those reactions without ascorbic acid, substrate inhibition was also considered while fitting the data in Michaelis-Menten equation of GraphPad Prism8 based on the default best-fit model. Further, the physiologically preferred substrate of BrDOD1 enzyme was determined using the stead-state kinetic data with the equation (Eq. 1) (Eisenthal et al., 2007), and the transition point substrate concentration ([S]_=_) above which both *K*_M_ and *V*_max_ increased was also determined using equation (Eq. 2) (Silverstein, 2019).

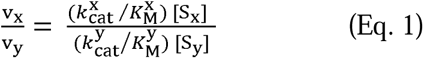

Where, v_x_ and v_y_ (velocity of substrate x and y) 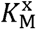 and 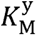 (Michealis-menten constant of substrate x and y); 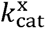 and 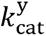 (turnover number towards substrate x and y); S_x_ and S_y_ (concentration of substrate x and y).

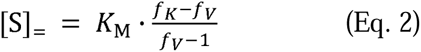

Where, [S]_=_ (Transition substrate point); *K*_M_ (Michealis-menten constant); *f*_K_ and *f_V_* (fold increase in *K*_M_ and *V*_max_).

### 2.8. Structure modelling, docking, and MD simulation

3D-structure of BrDOD1 protein was modelled followed by validations and then, molecular docking was performed. The protein ligand complex was used for molecular dynamics simulation of the complex on Schrodinger Maestro software (Academic version) (detailed protocol in the supplementary method file section 1.3.).

### 2.9. Mining of other LigB homologs

As it is already known that there are more than one LigB homologs in betalainic plants (Sheehan et al., 2020), to isolate the other paralogs, primers (Table S1A) for partial gene sequence amplifications were designed using the IDT OligoAnalyzer tool. The partial gene sequences of the other two homologs were amplified using *B. alba* L. var. ‘Rubra’ cDNA under optimized PCR conditions. The PCR products were purified using the GeneJET PCR purification kit and subjected to Sanger sequencing, and validated as mentioned above. Pair-end transcriptome data of *B. alba* (SRA: ERX2099282) were retrieved from the NCBI SRA database, and the transcriptome was subsequently trimmed and assembled using the Galaxy server (https://usegalaxy.org/). Using the standalone BLAST program of BioEdit 7.7.1, the partial sequences of the two homologs were searched against the assembled transcripts of *B. alba*, and the putative full-length sequences were retrieved. Primers (Table S1C) were then designed based on these sequences to amplify the full-length genes under optimized conditions. Cloning and confirmation of putative full-length LigB homologs were conducted as described above.

### 2.10. Phylogenetic tree and in silico analysis of LigB homologs

Further using the LigB homolog sequences of *B. alba* L. var. ‘Rubra’ or LigB homolog sequences of the order Caryophyllales, secondary structural alignment, group-specific conserved motif regions, and pI distribution were also generated (detailed protocol in the supplementary method file section 1.4., 1.5., 1.6, and 1.7).

## 3. Results

### 3.1. L-DOPA is the physiological substrate of BrDOD1

Partial sequence of a LigB homolog having length of 223 bp was amplified from the cDNA of *B. alba* L. var. ‘Rubra’ using gene-specific primers. Then, through RACE-PCR, the unknown flanking regions of its 5′ and 3′ ends were amplified and validated to have 176 bp and 912 bp, respectively (Fig. S3A). The RACE-PCR product sequences were assembled into a putative full-length gene of 981 bp (Fig. S3B). Further, using the open reading frame (ORF) specific primer, a full-length gene of 813 bp was amplified (Fig. S3A). This gene was named *BrDOD1*. After ligation to the cloning vector, the ligation was confirmed through restriction digestion (Fig. 1A) and then, and further ligated to expression vector followed by mobilization into *E. coli* BL21-CodonPlus cells. Isopropyl thiogalactoside (IPTG)-induced recombinant cells growing in M9 medium containing ampicillin and chloramphenicol carrying pLATE51:*BrDOD1* showed rapid formation of greenish yellow (λ_max_ 430 nm) after addition of substrate L-DOPA in the culture medium (Fig. S4). Further, L-DOPA dose-dependent response in the color formation was observed (Fig. 1B), thereby confirming the successful expression of BrDOD1 in *E. coli* BL21-CodonPlus cells. Recombinant BrDOD1 was extracted, purified (Table 1A) to homogeneity using Ni-NTA resin, desalted, and resolved on SDS-PAGE, and found to have a molecular weight of ∼30 kDa (Fig. 1C). Activity assays in different pH range after the Ni-NTA purification and enzyme activation showed that BrDOD1 showed the maximum activity in pH 6.8 (Fig. 1D). Further, enzyme assays using the purified protein showed the formation of greenish yellow (λ_max_ 430 nm) in the presence of L-DOPA or dopamine, as revealed by wavelength scan of the reaction product using a spectrophotometer. High performance liquid chromatography (HPLC) analysis of the reaction products BrDOD1 in the presence of substrate L-DOPA showed that BA eluted at a retention time of 10.6 min (λ_max_406 nm) and dopaxanthin at 9.25 min (λ_max_ 471 nm) (Fig. 1E), whereas dopamine-derived 6-decarboxy-BA eluted at a retention time of 13.1 min (λ_max_ 416 nm) and 6-decarboxy betaxanthin at 11.14 min (λ_max_ 470 nm) (Fig. 1F). The chromatogram profile was similar to that of previously published CqDOD2 reaction products and the identity of the peaks were confirmed through comparison (Henarejos-Escudero et al., 2022).

**Table 1A.** Fold purification of *Basella alba* L. var. ‘Rubra’ DOD1.

| Purification step | Total volume (mL) | Total proteins (mg) | Proteins (mg mL<sup>-1</sup>) | Activity (nmol min<sup>-1</sup> mL<sup>-1</sup>) | Total activity (nmol min<sup>-1</sup>) | Specific activity (nmol min<sup>-1</sup> mg<sup>-1</sup>) | Yield (%) | Fold purification |
| --- | --- | --- | --- | --- | --- | --- | --- | --- |
| Crude extract | 38.62 | 5319.6 | 137.7 | 26.1 | 1008.8 | 0.19 | 100 | 1 |
| Affinity (Ni-NTA) | 9.3 | 11.4 | 1.2 | 9.1 | 85.2 | 7.5 | 8.4 | 39.4 |
| Desalting (PD-10) | 12.6 | 9.8 | 1.04 | 25.8 | 325.6 | 33.2 | 33.2 | 175.2 |

**Fig. 1.**
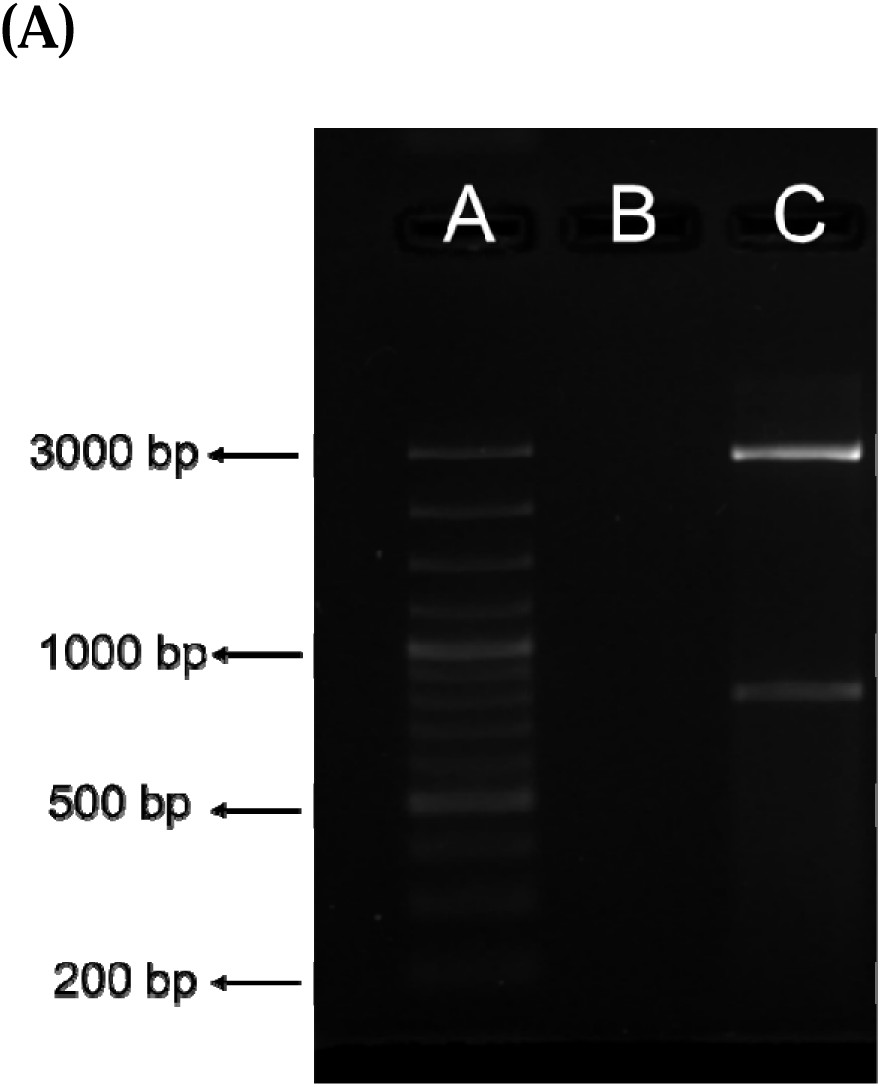
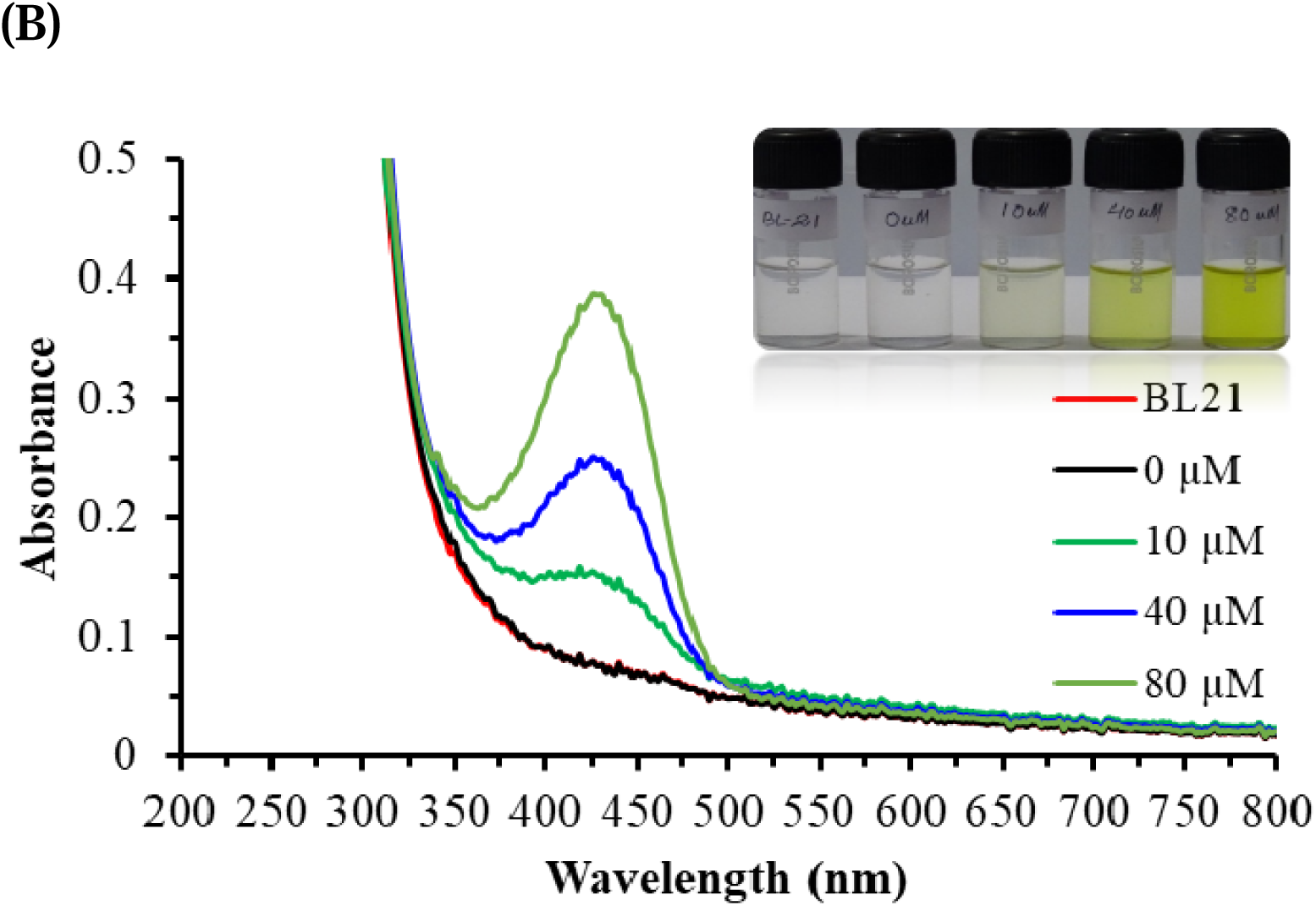
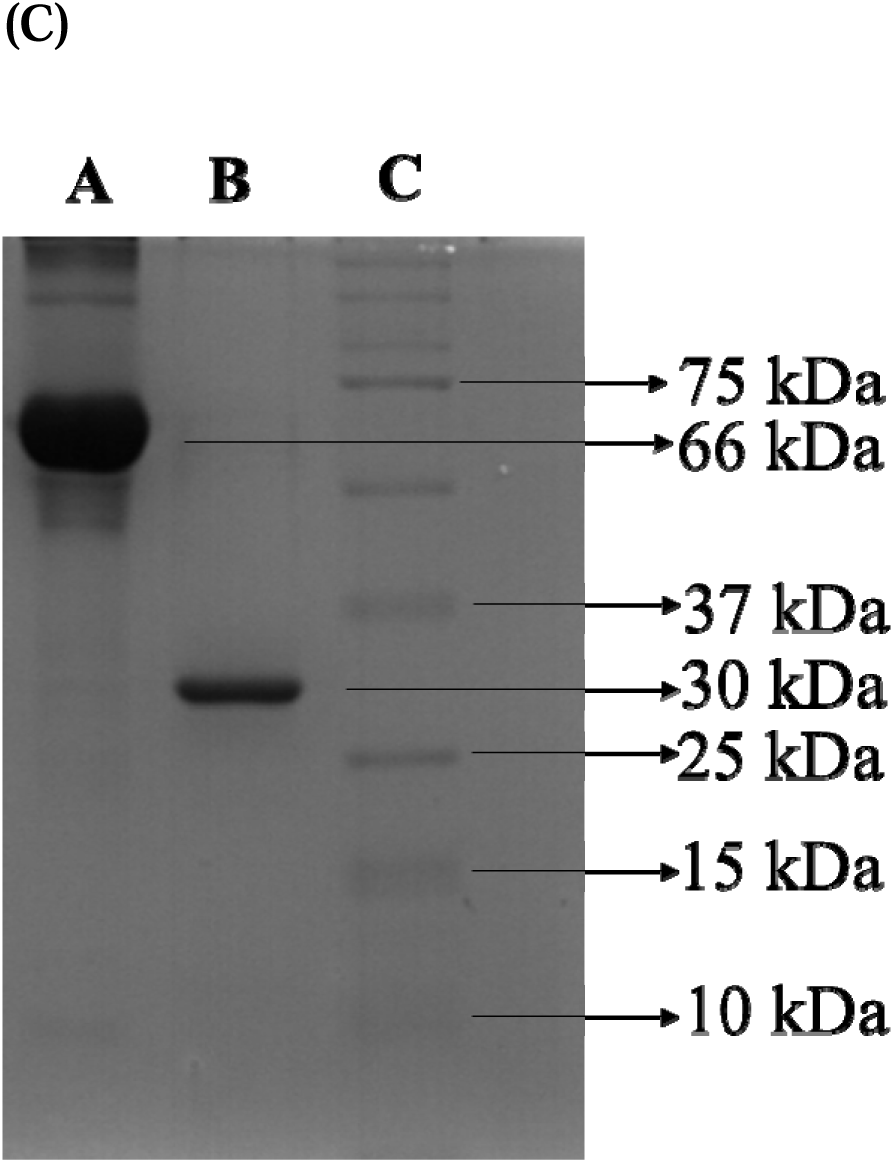
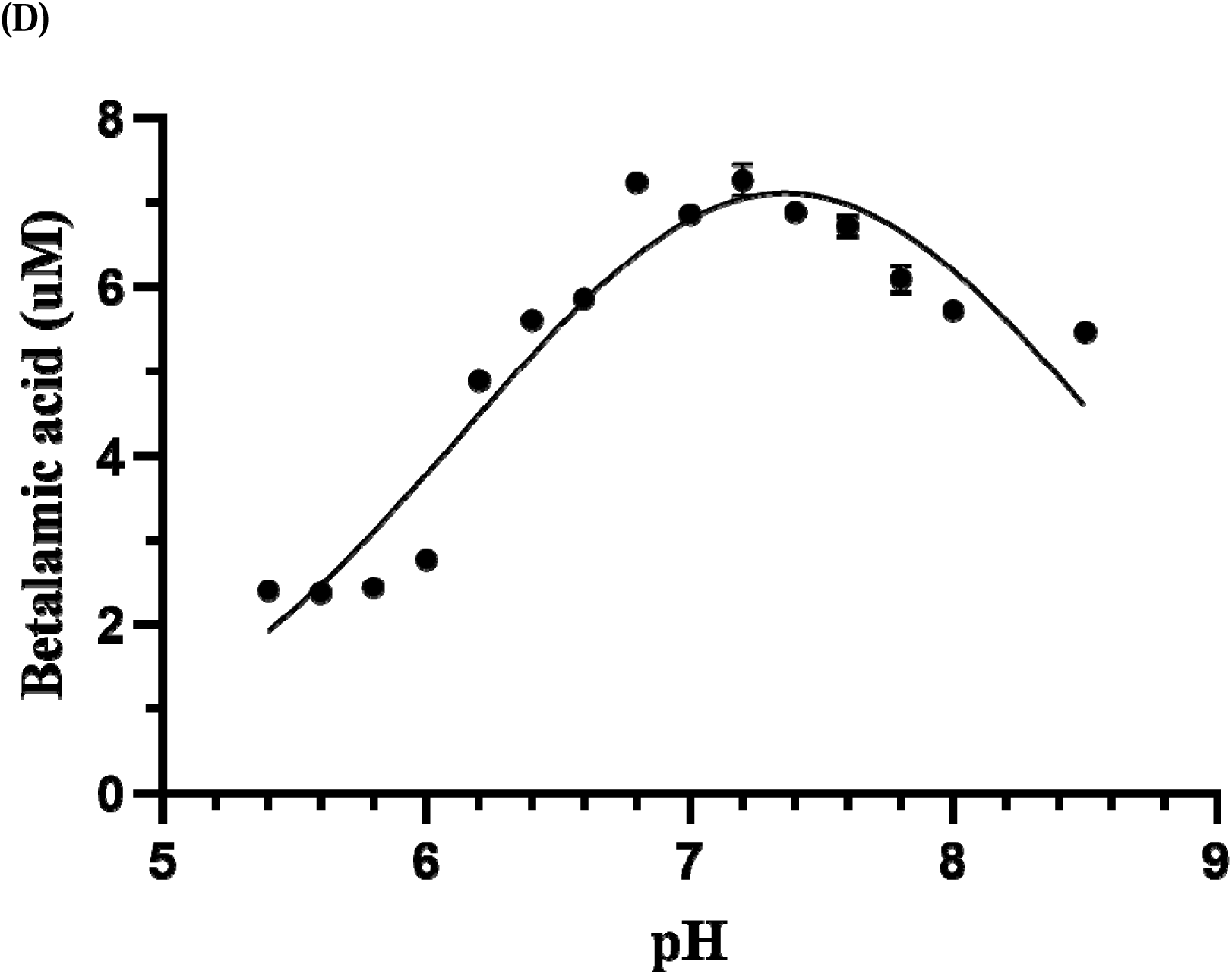
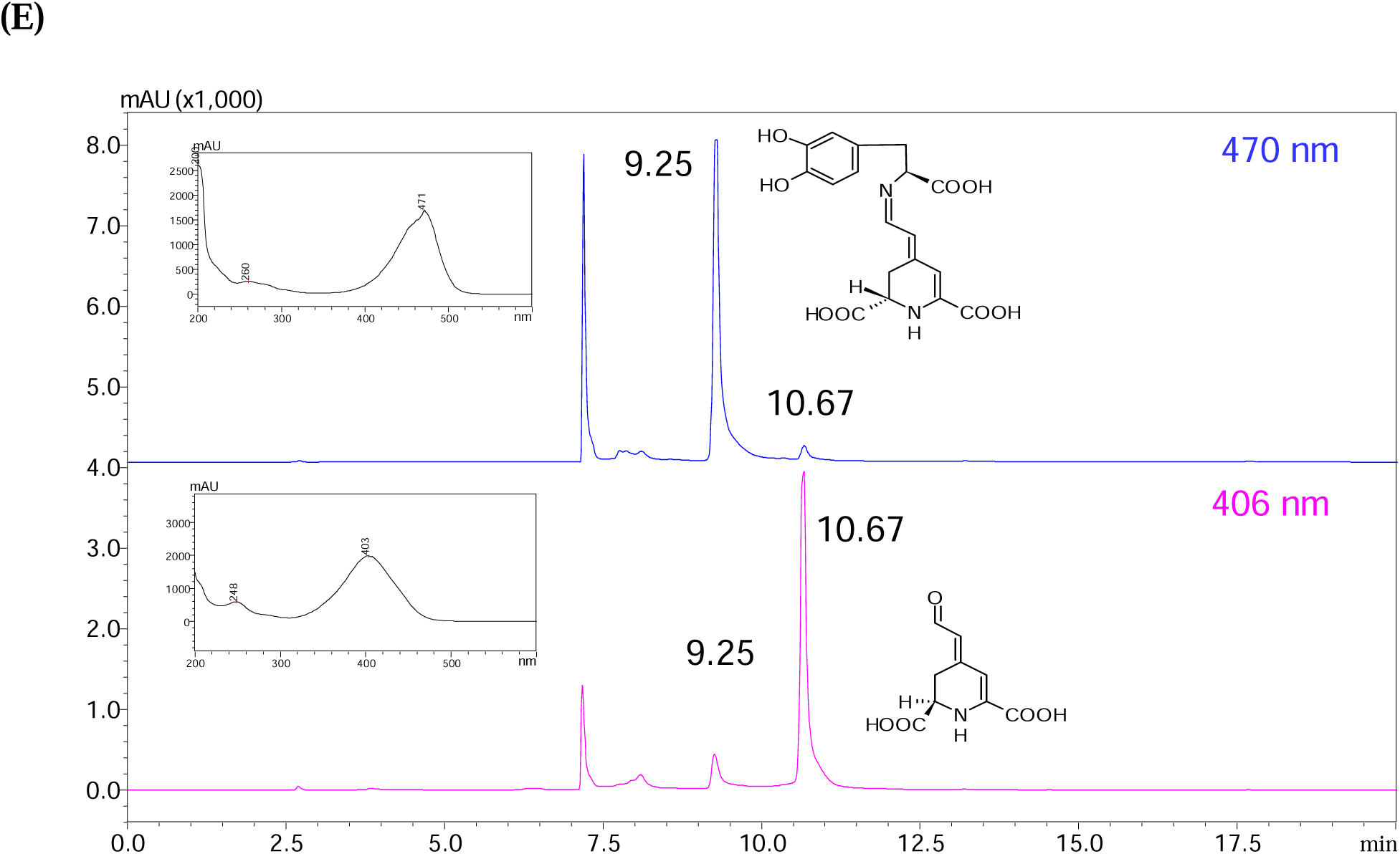
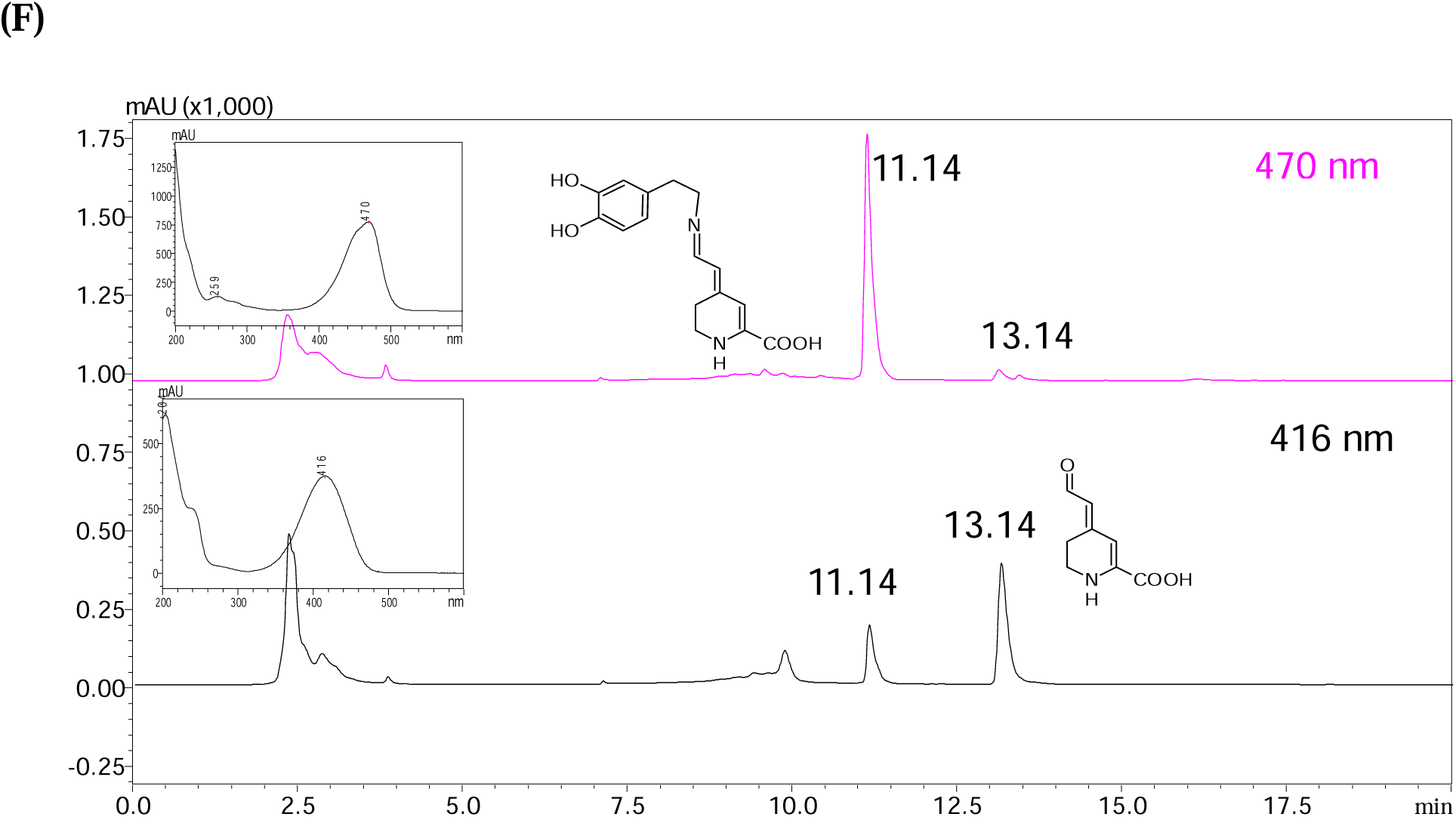
Heterologous expression and functional characterization of BrDOD1. **A)** Gel image showing restriction digested plasmid DNA (pJET1.2::*BrDOD1*) after transformation to *E. coli* DH5α cells (A- DNA ladder; B- blank well; C- pJET1.2::*BrDOD1* where the small band corresponds to *BrDOD1* while the larger one corresponds to digested plasmid), **B)** graph showing the betalamic acid forming-activity of recombinant pLATE51::BrDOD1 in the presence of different L-DOPA concentrations (**inset:** color difference of M9 medium after betalamic acid formation in the presence of different concentrations of L-DOPA by transformed *E. coli* BL21-CodonPlus cells), **C)** SDS-page gel picture showing homogeneity of purified BrDOD1 protein after Ni-NTA affinity chromatography (A- BSA; B- BrDOD1; C- molecular weight markers), **D)** non-linear curve fit showing the effect of different pH (50 mM) of acetate (4.0 – 5.6), phosphate (5.8 – 7.8), and tris-HCl (8.0 - 8.5) buffers at 25±2 °C on the betalamic acid-forming activity of BrDOD1, **E)** HPLC chromatograms showing detection of betalamic acid peak at 10.67 min and L-DOPA-derived betaxanthin peak at 9.25 min (**inset:** Absorption spectra of the two peaks), **F)** HPLC chromatograms showing detection of 6-decarboxy betalamic acid peak at 13.1 min and dopamine-derived betaxanthin at 11.4 min (**inset:** Absorption spectra of the two peaks).

Determination of steady-state kinetics in the presence of 10 mM ascorbic acid through non-linear regression showed that BrDOD1 had a *K*_M_ of 203.9 ± 6.2 *µ*M and a *V*_max_ of 5.9 ± 0.12 *µ*M min^-1^ toward L-DOPA (Fig. 2A; Table 1B), whereas *K*_M_ and *V*_max_ values toward dopamine were 360 ± 10.7 *µ*M and 1.9 ± 0.05 *µ*M min^-1^, respectively (Fig. 2B; Table 1B). Also, the linear steady-state kinetic parameters for both L-DOPA and dopamine were more or less similar with that of the non-linear regression analysis (Table S2; Fig. S5A and B). However, in the absence of ascorbic acid, the kinetic parameters were greatly affected. Without ascorbic acid, BrDOD1’s affinity was much higher (*K*_M_ of 31.0 ± 0.81 *µ*M for L- DOPA and 46.5 ± 2.9 *µ*M for dopamine). The *V*_max_ values too were greatly reduced to 1.7 ± 0.02 *µ*M min^-1^ and 0.32 ± 0.01 *µ*M min^-1^ toward L-DOPA and dopamine, respectively (Fig. 2C and D; Table 1B). Further, in the absence of ascorbic acid, substrate inhibition was observed in both substrates with an inhibition constant (*K*_i_) of 7552.5 ± 552.8 *µ*M for L- DOPA and 3493 ± 367.5 *µ*M for dopamine. Based on the reaction rate toward L-DOPA (*v*_DOPA_or *v_x_*) and dopamine (*v*_dopamine_ or v*_y_*), the ratio *v*_DOPA_/*v*_dopamine_ was 6.6, whereas the ratio of *v*_dopamine_/*v*_DOPA_ was 0.15 throughout the tested substrate concentrations (12.5 – 3200 *µ*M), indicating that BrDOD1’s preferred substrate could be L-DOPA because of the 6.6-fold higher reaction rate toward it than that of dopamine. In view of this observation, we analyzed the molar concentrations of L-DOPA and dopamine in *B. alba* L. var. ‘Rubra’ fruits using HPLC. The L-DOPA content was significantly higher than that of dopamine (Table 2; Table S3) in mature/ripe fruit pulp, indicating that BrDOD1 might prefer L-DOPA physiologically because of its higher content than dopamine.

**Table 1B.** Steady-state kinetics of *Basella alba* L. var. ‘Rubra’ DOD1 in the presence or absence of ascorbic acid.

| Assay condition | $V_{\max}$ ($\mu\text{M min}^{-1}$) | $K_M$ ($\mu\text{M}$) | $K_i$ ($\mu\text{M}$) | $k_{\text{cat}}$ ($\text{min}^{-1}$) | $k_{\text{cat}}/K_M$ ($\text{nM}^{-1} \text{min}^{-1}$) |
| --- | --- | --- | --- | --- | --- |
| L-DOPA | $1.7 \pm 0.02$ | $31.0 \pm 0.81$ | $7552.5 \pm 552.8$ | $0.57 \pm 0.008$ | $18.5 \pm 0.53$ |
| Dopamine | $0.26 \pm 0.01$ | $46.5 \pm 2.9$ | $3493 \pm 367.5$ | $0.09 \pm 0.0003$ | $1.9 \pm 0.09$ |
| L-DOPA with AscA | $5.9 \pm 0.12$ | $203.9 \pm 6.2$ | NA | $2.01 \pm 0.03$ | $9.9 \pm 0.2$ |
| Dopamine with AscA | $1.6 \pm 0.04$ | $360.4 \pm 10.7$ | NA | $0.54 \pm 0.01$ | $1.5 \pm 0.03$ |

AscA- ascorbic acid; Mean ± SD (n=4)

**Fig. 2.**
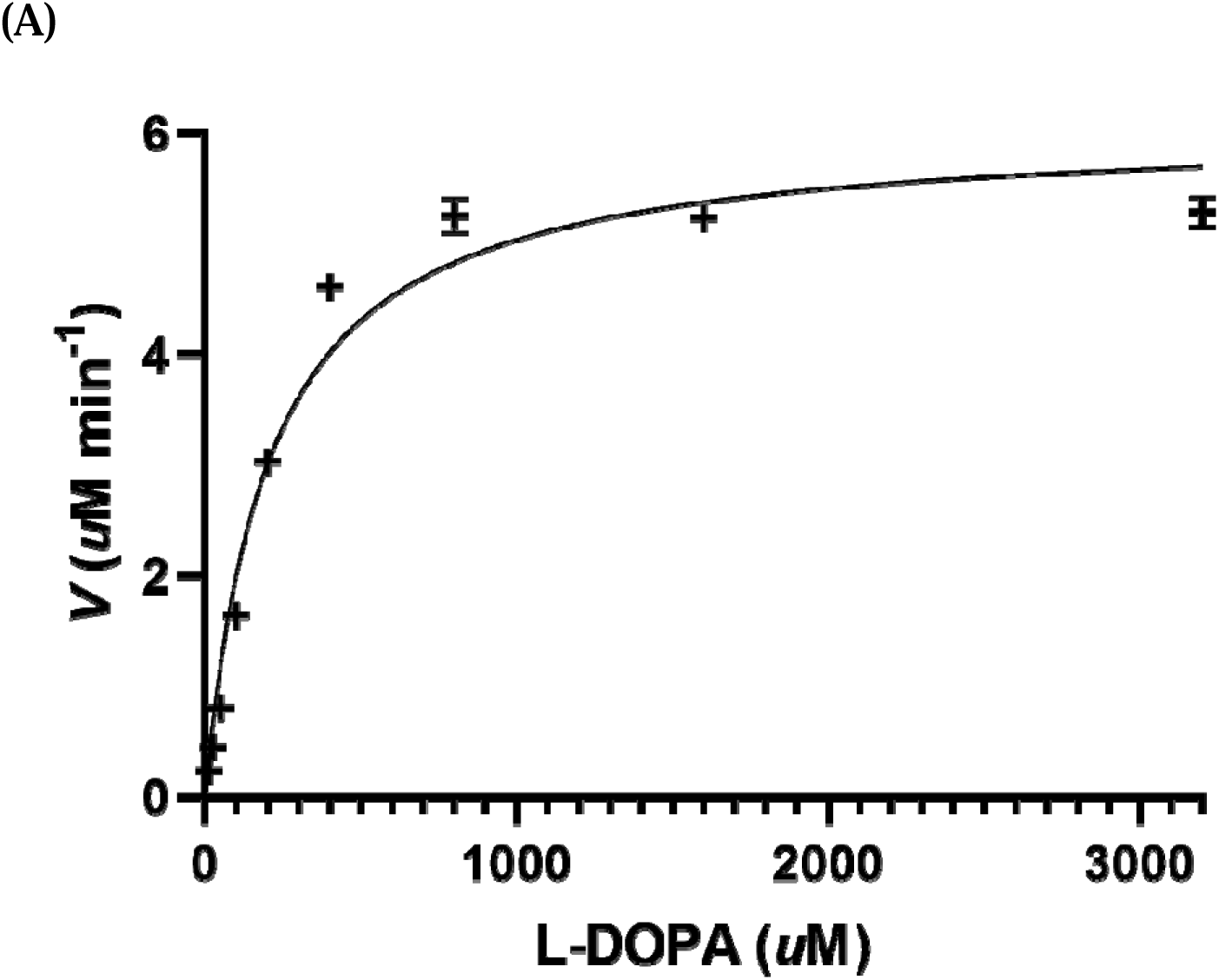
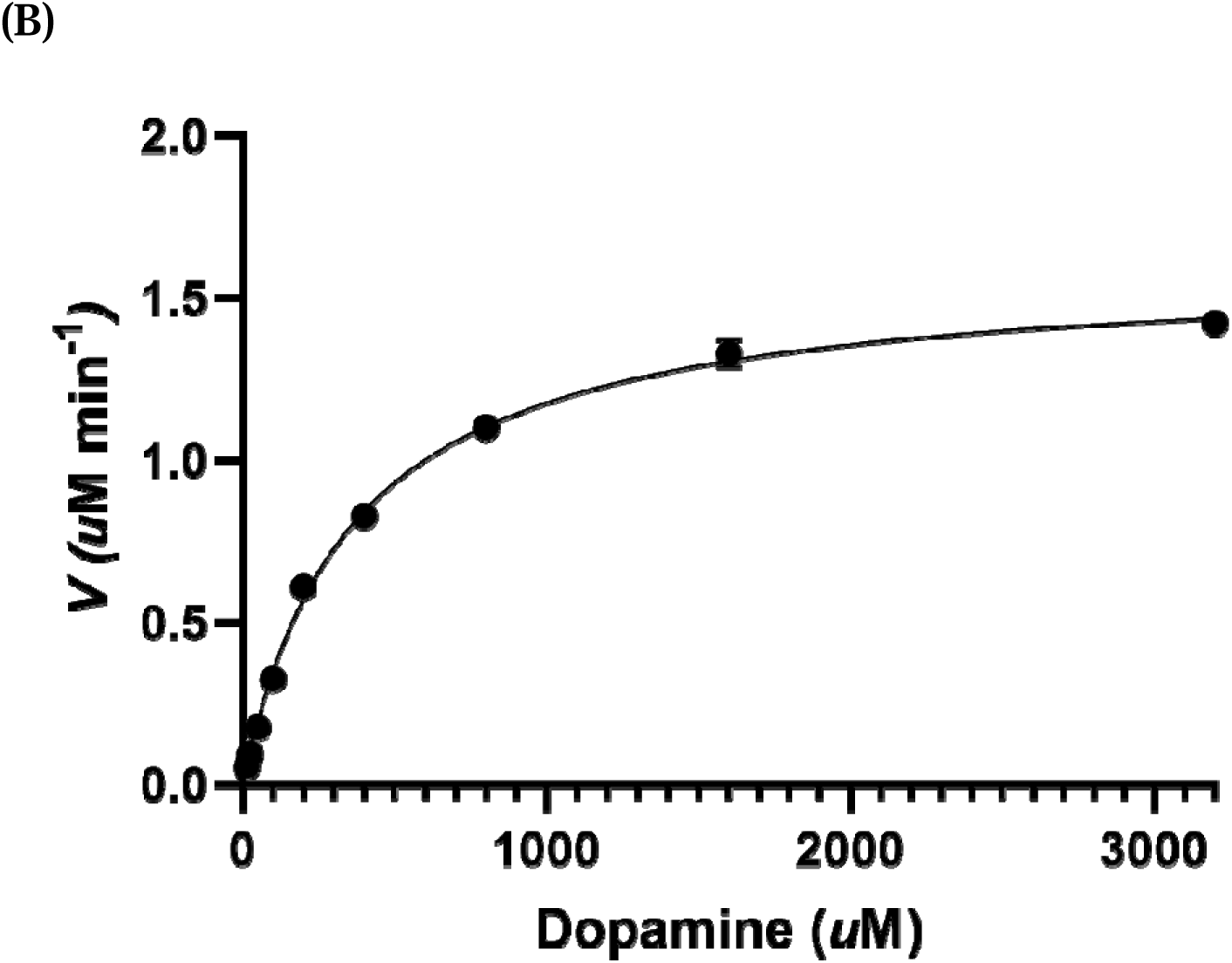
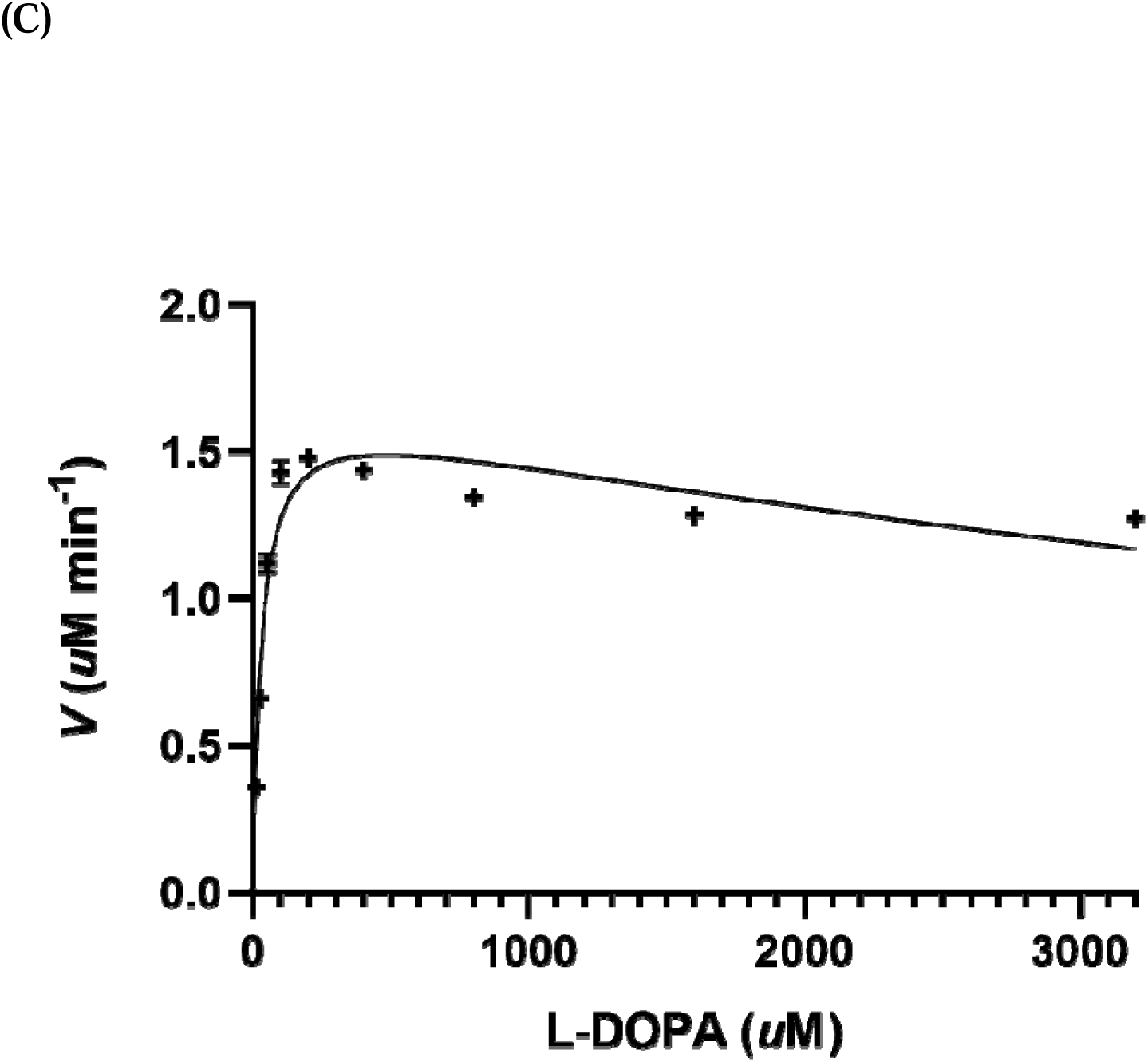
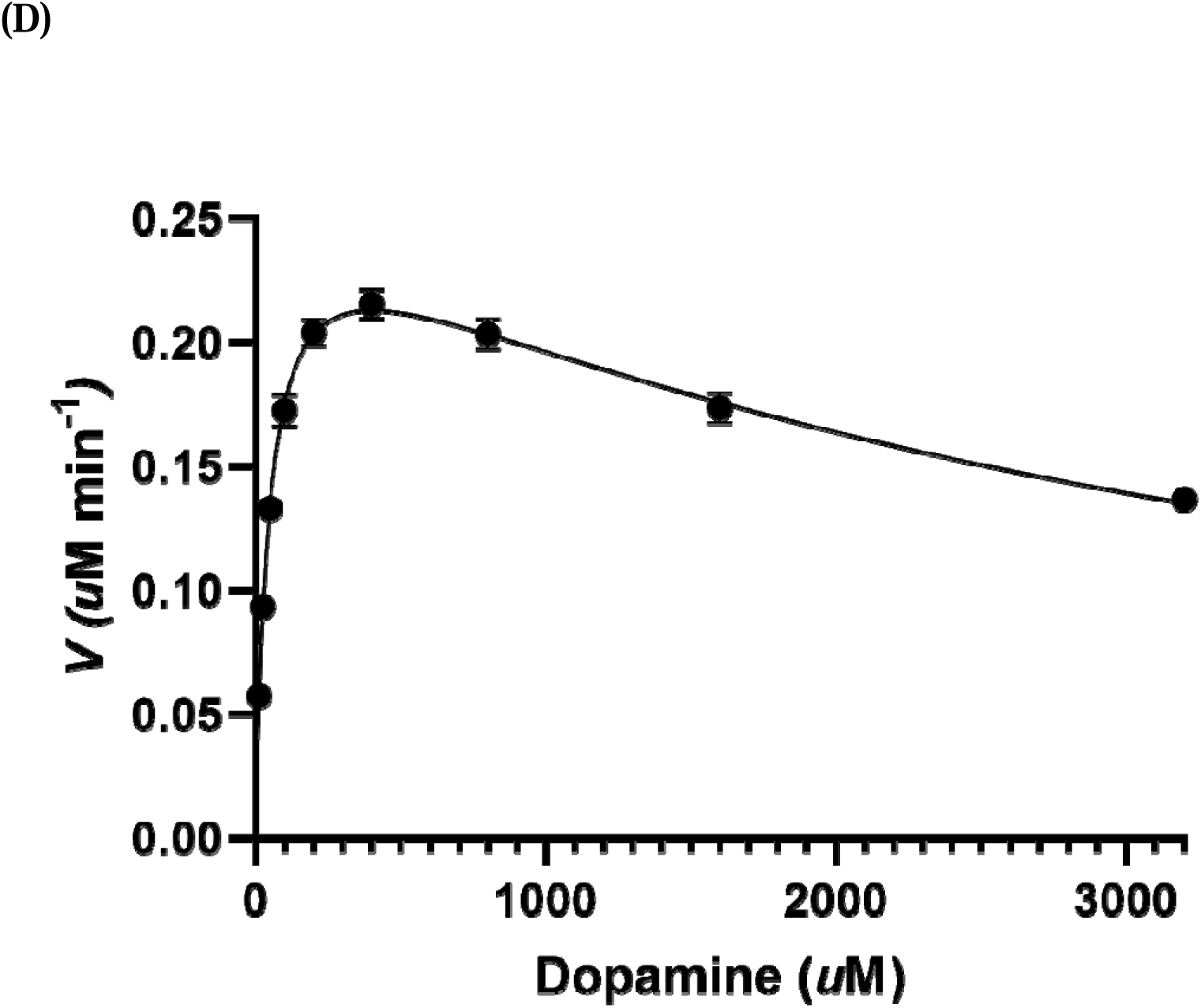
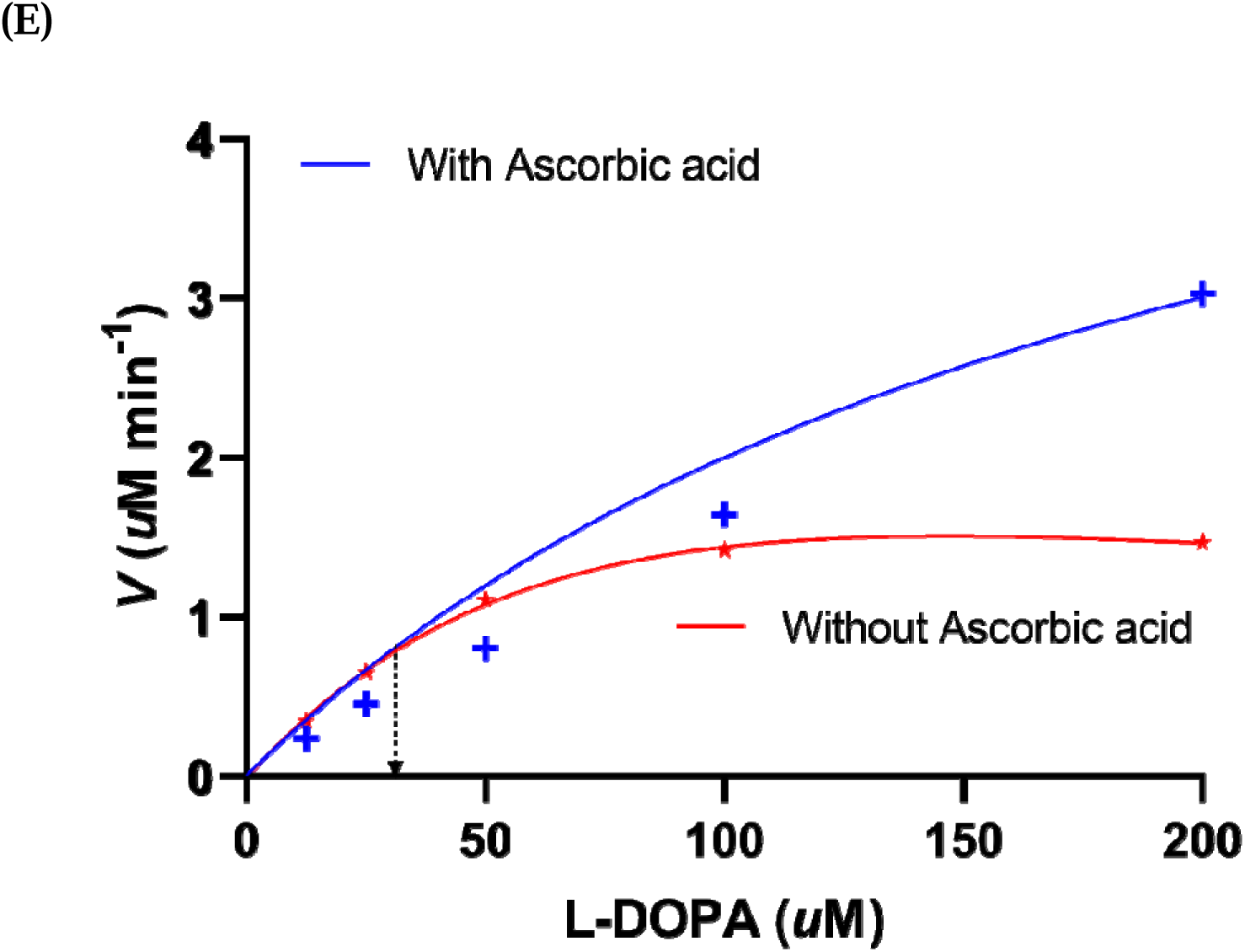
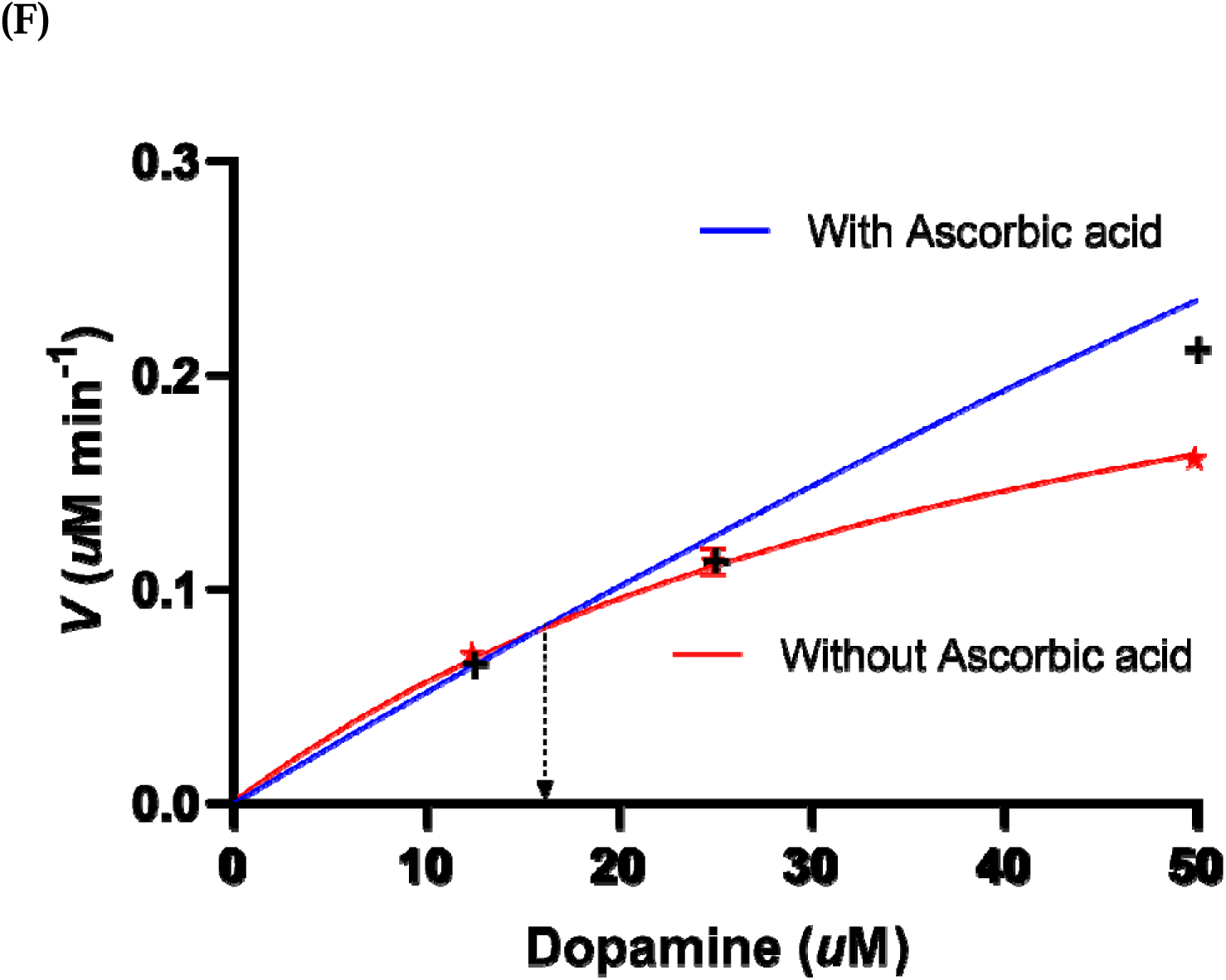
Catalytic kinetics of BrDOD1 using non-linear regression. Steady-state Michaelis-Menten kinetics of BrDOD1 in the presence of **A)** L-DOPA and ascorbic acid, **B)** dopamine and ascorbic acid, **C)** L-DOPA without ascorbic acid, **D)** dopamine without ascorbic acid, **E)** overlay of steady-state Michaelis-Menten kinetic curves in the presence of L-DOPA with or without ascorbic acid showing transition substrate concentration, **F)** overlay of steady-state Michaelis-Menten kinetic curves in the presence of dopamine with or without ascorbic acid showing the transition substrate concentration. All analyses were performed using GraphPad prism8 using default best-fit model.

**Table 2.** Molar concentrations* of L-DOPA and dopamine in betalainic plants based on fresh weight and tissue moisture content.

| Sample | L-DOPA<br>($\mu\text{M g}^{-1}$) | Dopamine<br>($\mu\text{M g}^{-1}$) | Ascorbic acid<br>($\mu\text{M g}^{-1}$) | Total betalain<br>content (mg g<sup>-1</sup>) | Reference |
| --- | --- | --- | --- | --- | --- |
| *Basella alba* L. var.<br>“Rubra” fruit<br>(mature/ripe pulp) | 727.3 ± 9.8 | 398.1 ± 23.7 | 2393.6 ± 62.3 | 1.2 ± 0.03 | This study |
| *Amaranthus tricolor*<br>red seedlings (12-day-old) | 51.9 ± 2.4 | 8.0 ± 0.77 | 1083.1 ± 59.2 | 0.39 ± 0.02 | (Kumari et al., 2024) |
| *Amaranthus tricolor*<br>red seedlings (16-day-old) | 21.2 ± 0.48 | 9.9 ± 0.53 | 748.3 ± 51.7 | 0.36 ± 0.007 | |
| *Amaranthus tricolor*<br>seedlings (30-day-old) | 1.2 ± 0.06 | 0.94 ± 0.09 | 2557 ± 101.9 | 0.22 ± 0.02 | |
| *Celosia argentea* var.<br>cristata (L.) Kuntze<br>inflorescence (yellow) | nr | 41.1 | Not studied | 0.09 $\mu\text{mol}$ | (Schliemann et al., 2001) |

nr- not reported. Mean ± SEM (n=3 for *Basella* L. var. ‘Rubra’; 4 for *Amaranthus* L-DOPA and dopamine; 3 for *Amaranthus* ascorbic acid). The values are based on the fresh weight.
\* The actual physiological values will be higher considering that L-DOPA is present mostly in cytosol and most of the water content in plant cells is in vacuoles, which takes up volume $\geq 50\%$ .

### 3.2. Ascorbic acid induces molecular crowding effect on BrDOD1 activity

Table 1B shows that both *V*_max_ and *K*_M_ of BrDOD1 were increased in the presence of 10 mM ascorbic acid. To understand this phenomenon better, the two steady-state kinetic curves in the presence and absence of ascorbic acid were drawn together for each substrate. From this graph, the transition point substrate concentrations ([S]_=_) were observed to be 38.3 *µ*M and 14.5 *µ*M for L-DOPA and dopamine, respectively (Fig. 2E and F; Fig. S6A and B). Above this substrate concentration, ascorbic acid (10 mM) acted as an activity enhancer, whereas below this substrate concentration, the effect was inhibitory. Such an effect of cosolute (ascorbic acid, in this case) included in much higher concentration in the assay compared to that of substrate and enzyme, may be due to favoring of association reactions of the protein and substrate against dissociation reaction as seen in the case of small molecule crowders (Silverstein, 2019). Physiologically, the *B*. *alba* L. var. ‘Rubra’ mature fruit pulp contained only 2393.6 ± 62.3 μM ascorbic acid/g fresh weight (Table 2; Table S3), indicating a very low concentration compared to the 10 mM ascorbic acid used in the enzyme assay. Even much lower concentration was that of L-DOPA (Table 2; Table S3). Considering the much higher physiological concentration of ascorbic acid than that of L-DOPA in the betalain-accumulating *B. alba* L. var. ‘Rubra’ fruit pulp (Table 2), it may be possible that ascorbic acid enhances the enzyme activity *in planta* as well. From Fig 2E and F, it is apparent that ascorbic acid at 10 mM produced crowding effect, thereby increasing both *K*_M_ and *V*_max_ of BrDOD1 toward L-DOPA as well as dopamine.

### 3.3. L-DOPA-BrDOD1 complex is more stable than dopamine-BrDOD1

Protein structure modelled on AlphaFold3 (Fig. S7A) was validated through Ramachandran plot, where 89.4% residues were mapped in the most favored region (Fig. S7B). Molecular docking of the substrate on the 3-D model of BrDOD1 revealed that L-DOPA showed a better binding (Δ*G* = - 5.7 kcal/mol) than dopamine (Δ*G* = - 4.9 kcal/mol). All LigB-characteristic residues (according to (Christinet et al., 2004)) His176, Asp179, and Asp180 interacted with dopamine, but L-DOPA interacted with His176 and Asp179 only among the three residues. In addition, Pro177 interacted with dopamine but not L-DOPA. The same Pro residue is diagnostic of the convergence of high BA-forming *Beta vulgaris* DODAα1 (BvDODAα1) from DODAβ (Brockington et al., 2015). Another proposed diagnostic residue Trp222 (Trp227 of BvDODAα1) did not interact with either L-DOPA or dopamine, indicating that the diagnostic residues may be involved in some other structural/functional role apart from substrate binding. In all, for the same cavity volume of 186 ^3^, L-DOPA interacted with 21 residues (His16, Gly17, Asn18, Pro19, Ala20, Phe27, His54, Asp73, Phe74, Ser75, Asp76, Val77, His120, His176, Asp179, Phe228, Asp230, His231, Asp256, Leu260, and Tyr262) (Fig. S7C and D), while dopamine did so with 32 residues (Ser15, His16, Gly17, Asn18, Pro19, Ala20, His54, Asp73, Phe74, Ser75, Asp76, Val77, His120, Gly172, Gly173, Ala174, Val175, His176, Pro177, Thr178, Asp179, Asp180, Thr181, Pro182, His183, His231, Asp256, His257, Thr259, Leu260, Gly261, and Tyr262) (Fig. S7E and F). Although dopamine interacted with more BrDOD1 residues compared to that of L-DOPA, the stability of the former was less than the latter, according to the root mean square deviation (RMSD) graphs obtained through molecular dynamics (MD) simulation for 100 ns (Figs 3A and B). Therefore, BrDOD1’s complex with L-DOPA (Fig. 3A) is more stable than its complex with dopamine (Fig. 3B), indicating that L-DOPA could be BRDOD1’s thermodynamically preferred substrate.

**Fig. 3.**
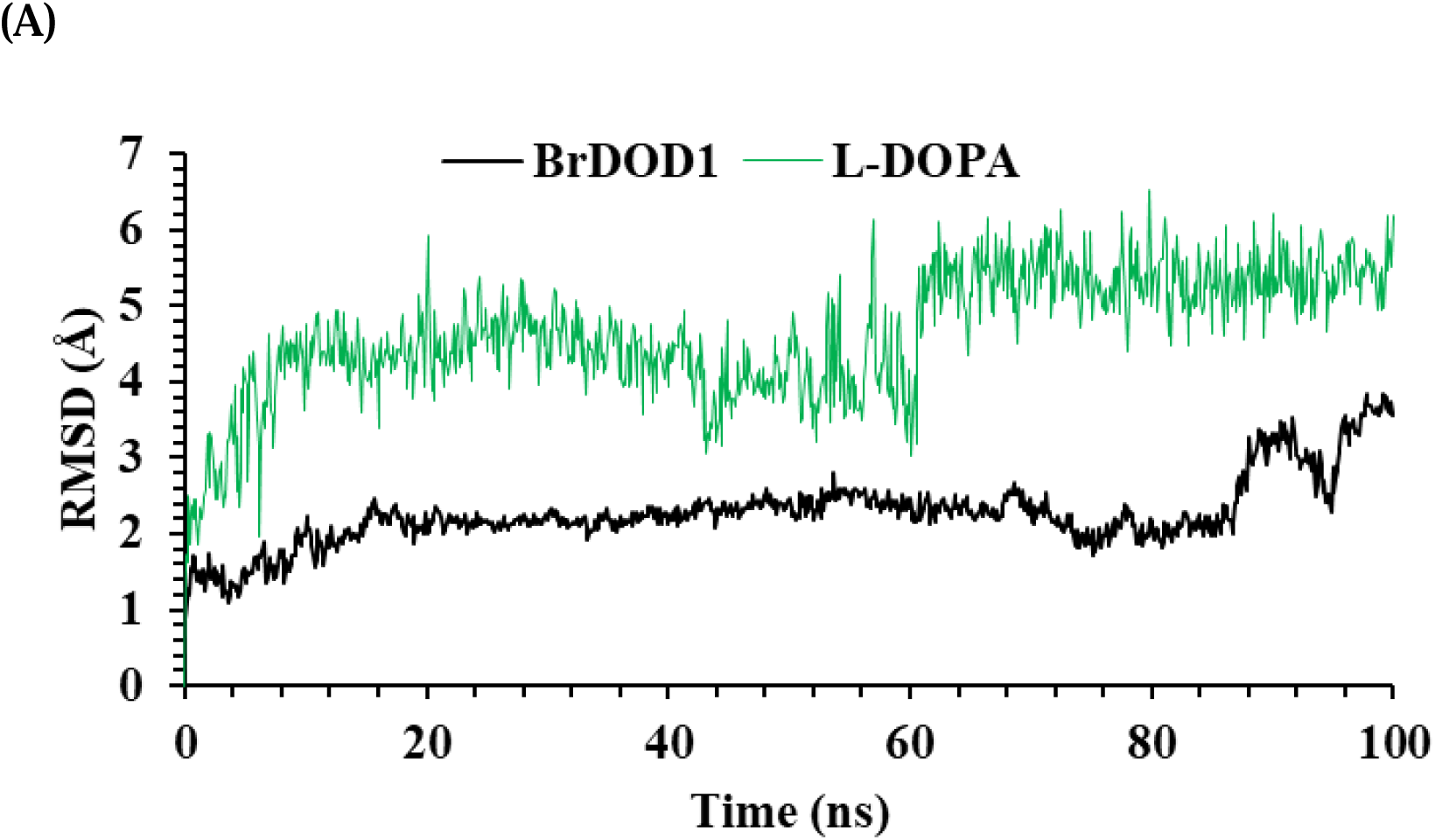
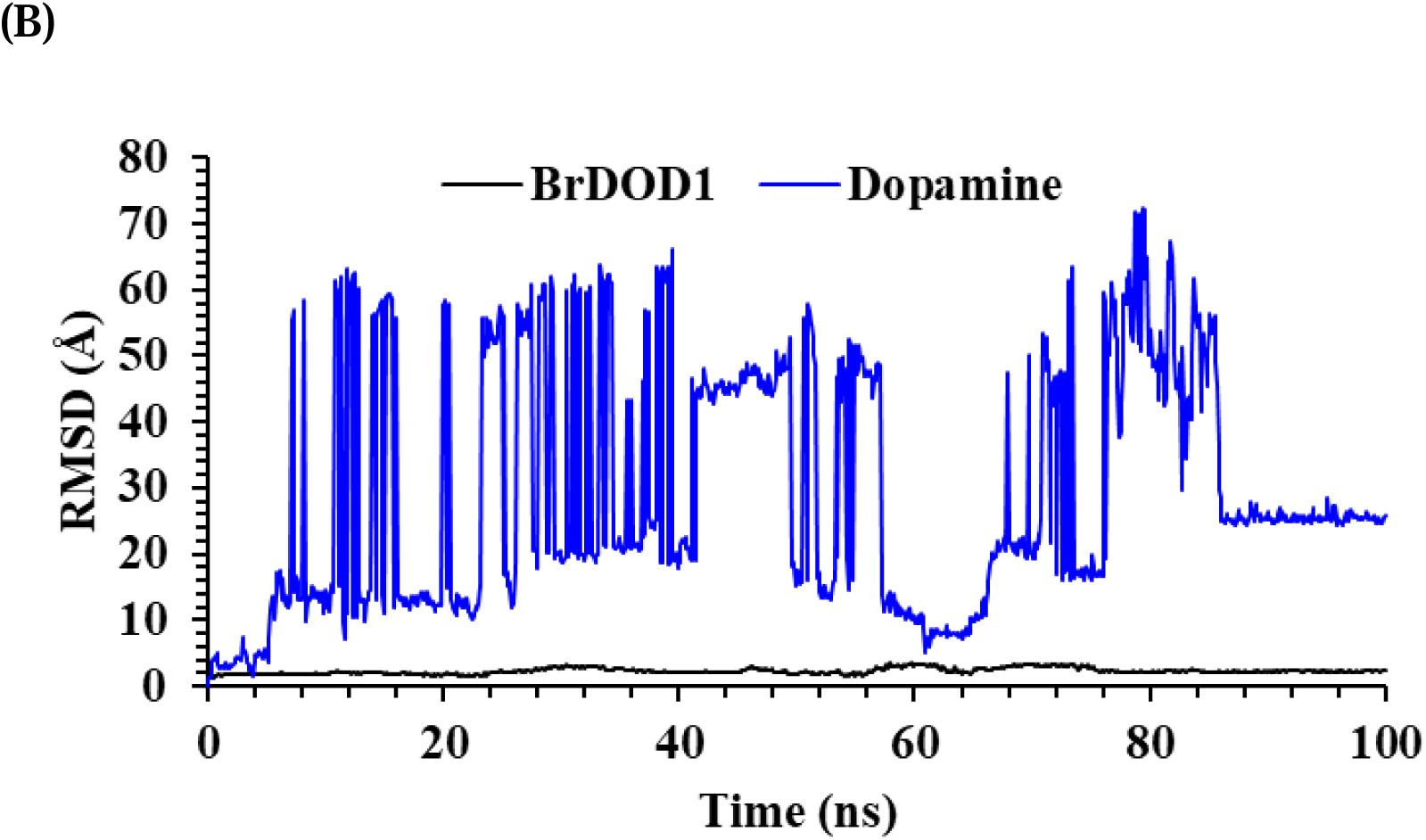
Molecular dynamics simulation of BrDOD1 with different substrates. RMSD plot of BrDOD1 docked with **A)** L-DOPA and **B)** dopamine showing more stability of the interaction with L-DOPA than that of dopamine over 100 ns.

### 3.4. *B. alba* L. var. ‘Rubra’ has three LigB homologs with distinctive features

Mining through the assembled transcriptome data using confirmed partial sequences of two more LigB homologs having 343 and 270 bp of *B. alba* L. var. ‘Rubra’, we found two putative LigB sequences. Through homology search on NCBI and ORFfinder tool, two putative full-length LigB homologs were identified. By using ORF specific primers, the full-length genes of the two LigB homologs having ORFs 792 bp and 996 bp were amplified and confirmed through sequencing (Fig. S8A). Further, analysis of sub-cellular targeting of all three LigB homologs isolated from *B. alba* L. var. ‘Rubra’ showed that BrDOD1 and the paralog of 792 bp (named BrDOD2) are targeted to both nucleus and cytoplasm, whereas the third homolog of 996 bp (named BrLigB) was targeted to endoplasmic reticulum (Table S5). The predicted pI values of these three full-length genes also varied (Table S5). Secondary structure analysis with respect to BvDODAα2 (PDB ID: 8IN2) revealed that all the three LigB homologs have distinctive conserved motifs near the active site forming a loop as part of the substrate-binding pocket (Chiang et al., 2025) (Fig. 4A). Further, phylogenetic tree of LigB homologs in betalainic plants under Caryophyllales showed three clades dividing the LigB homologs into LigB clade, which branches out into two more clades where DOD1 and DOD2 proteins were resolved (Fig. 4B). The two clades consisting of DOD1 and DOD2 proteins indicate the possible evolution of DOD1 from DOD2, as DOD2 appeared before DOD1 in both clades. Each one of the three LigB homologs of *B. alba* L. var. ‘Rubra’ fell in different clades (blue/LigB, red/DOD1, and green/DOD2), indicating clearly the phylogenetic divergence into three clades (Fig. 4B). BrDOD1 fell in the red clade alongside other high L-DOPA-derived BA-forming MjDOD1. BrDOD2 was monophyletic with MjDOD2 (green clade), which show low/marginal L-DOPA-derived BA-forming activity (Fig. 4B). BrLigB was ingroup with low/marginal L-DOPA-derived BA-forming proteins, including BvDODAβ (Brockington et al., 2015; Sheehan et al., 2020), which may be named BvLigB, colored in blue (Fig. 4B). Alignment of DOD1, DOD2, and LigB protein sequences across betalainic plants of Caryophyllales revealed that the conserved loop-forming sequences of the putative substrate-binding pocket (based on the crystal structure of BvDODAα2, PDB ID: 8IN2) of high BA-forming activity DOD1 group proteins had H-P-[S/L/T]-D-[D/E]-T-P, while that of low/marginal BA-forming activity DOD2 group proteins had H-P-N-[N/S/G]-T-P (Table 3). The H-N-L-R motif was present in the sequences of LigB group proteins only irrespective of their source, i.e., betalain-accumulating or betalain-devoid plants (Table 3 and Fig. S8B). The isoelectric points (pI) of DOD homologs (Fig. 4C) revealed that the median pI (5.67) of the high BA-forming DOD1 group was the lowest among the three groups, while low/marginal BA-forming DOD2 (pI 6.25) was in between, and low BA-forming LigB group showed the highest pI of 6.8.

**Fig. 4.**
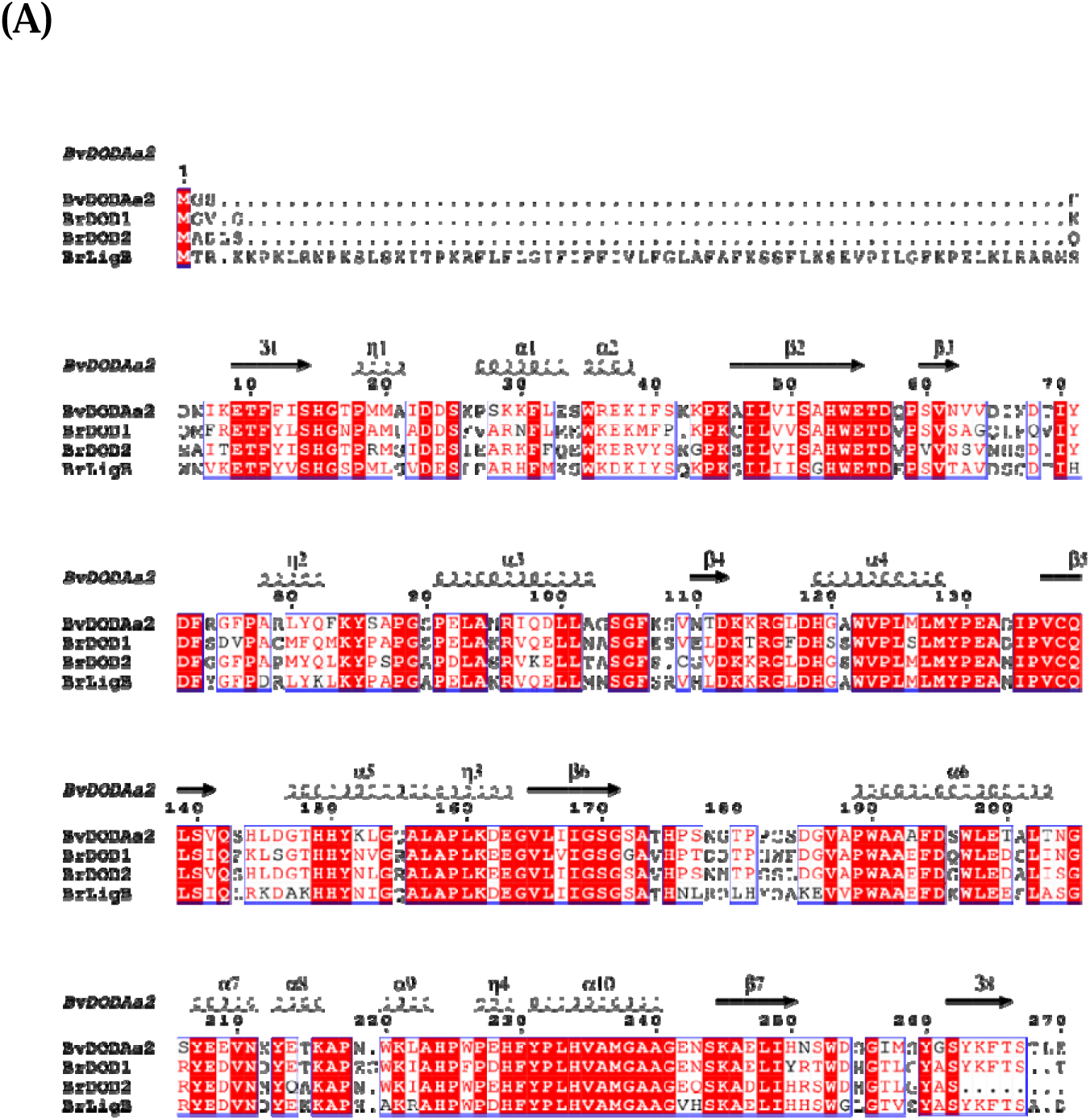
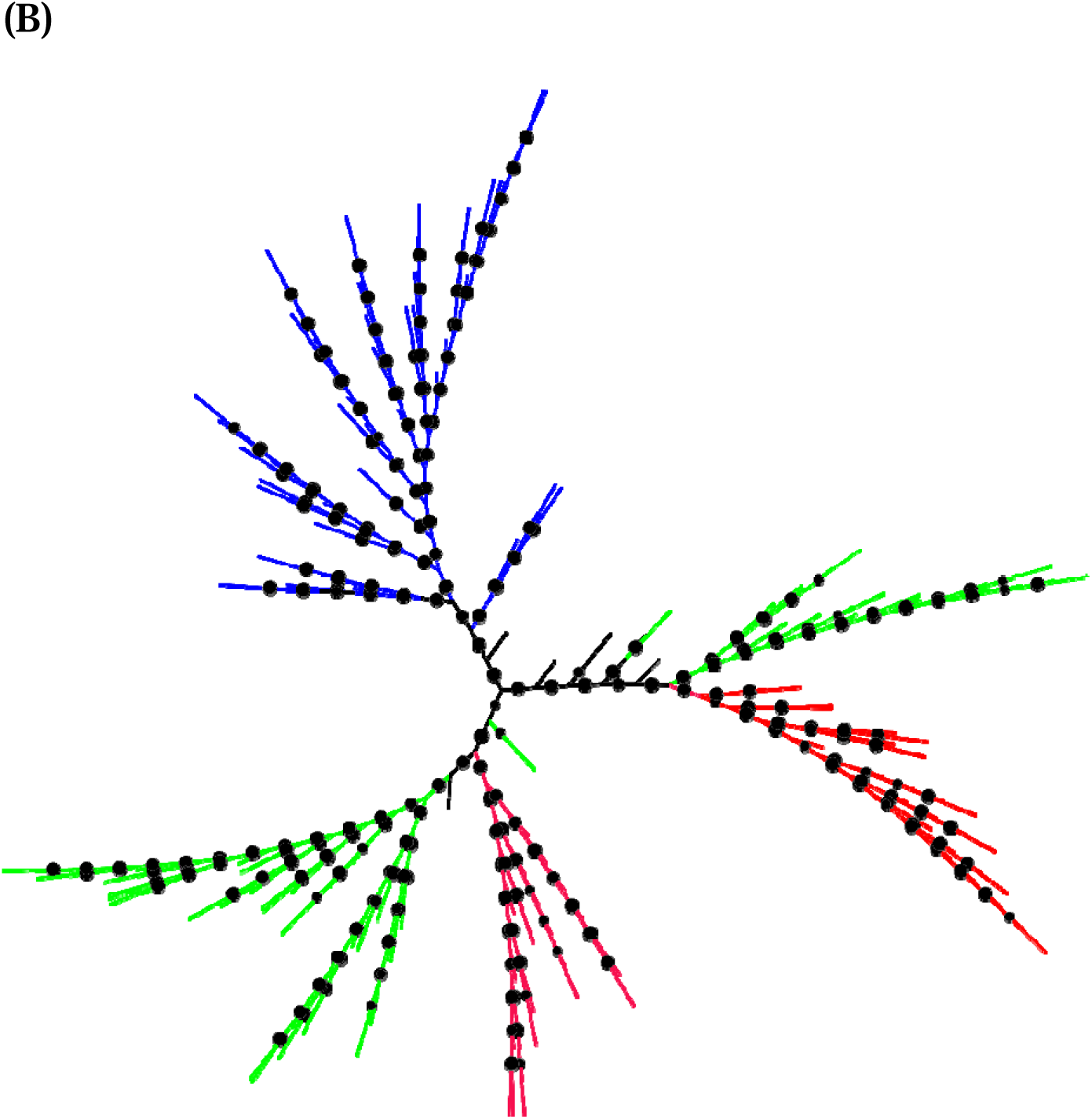
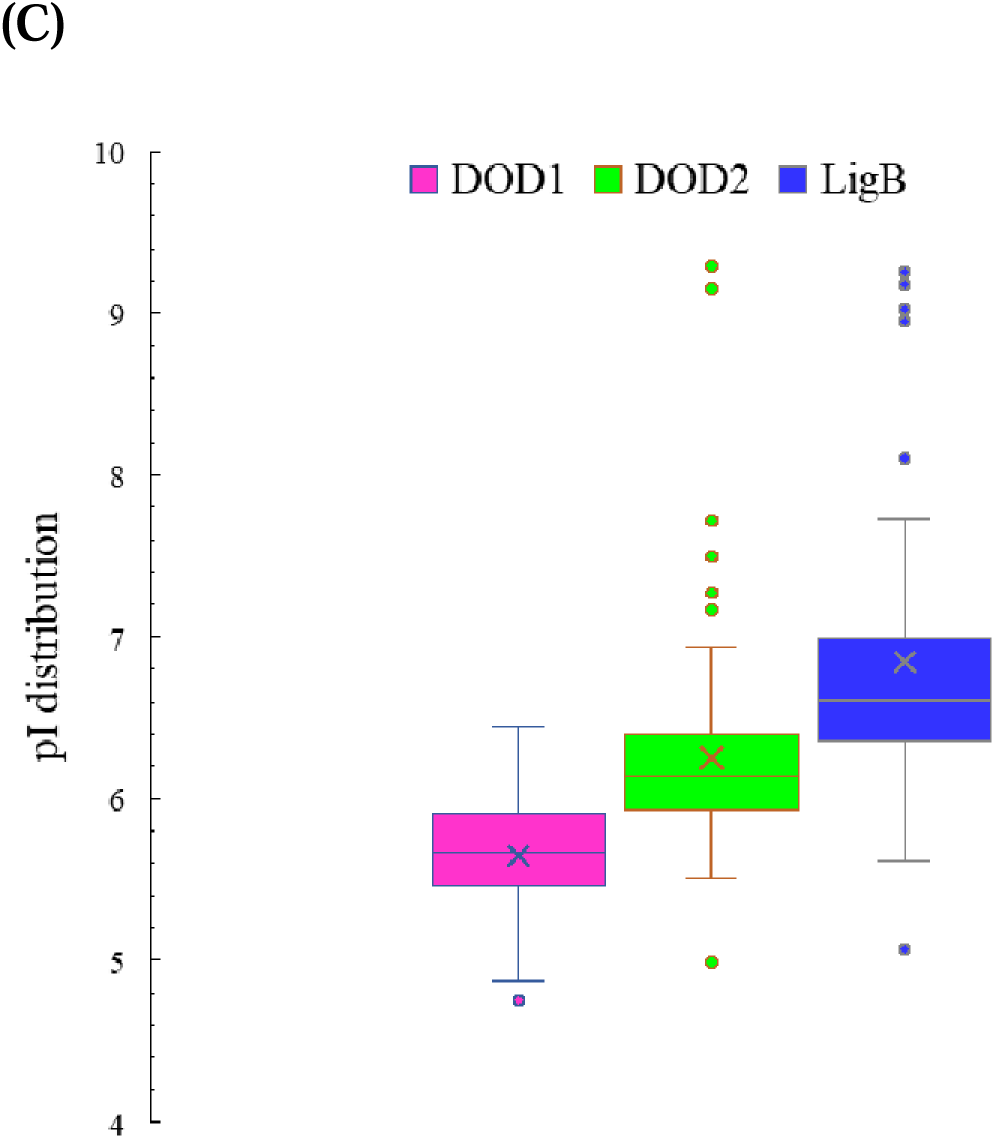
Characteristic features of LigB homolog sequences of *B. alba* L. var. ‘Rubra**’**. **A)** Secondary structure alignment showing the comparison and alignment of deduced BrDOD1, BrDOD2, and BrLigB protein sequences with beetroot DOD2 (BvDODAα2), the only plant DOD with a crystal structure (PDB ID: 8IN2), **B**) different classes of LigB homologs of plants under the order Caryophyllales (red-violet clade-high activity DOD1s, green clade-low/marginal activity DOD2s, and blue clade-LigB orthologs), and **C)** box-plot graph showing the distribution of deduced isoelectric point (pI) values of DOD1, DOD2 and LigB groups from the plant under the order Caryophyllales.

**Table 3.**
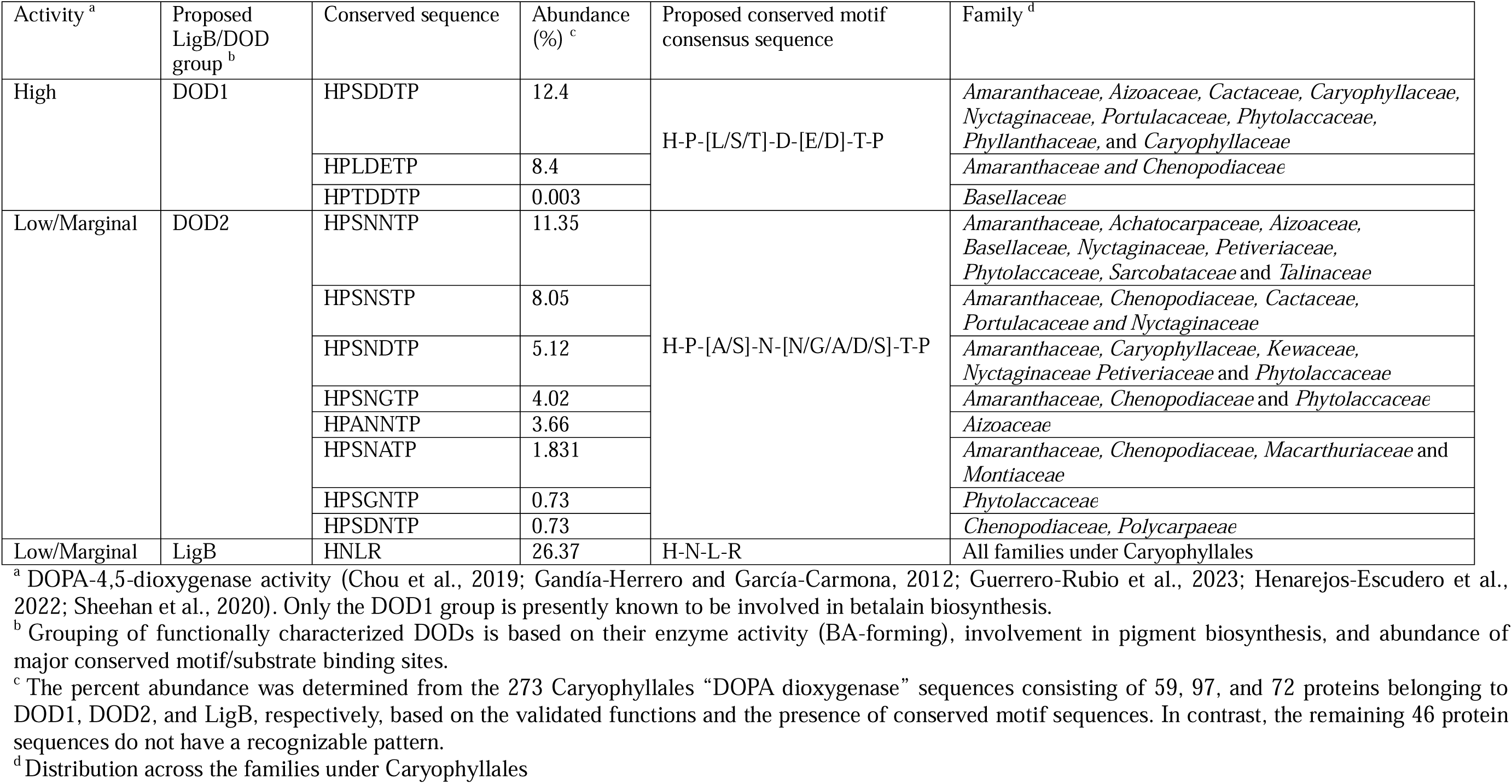
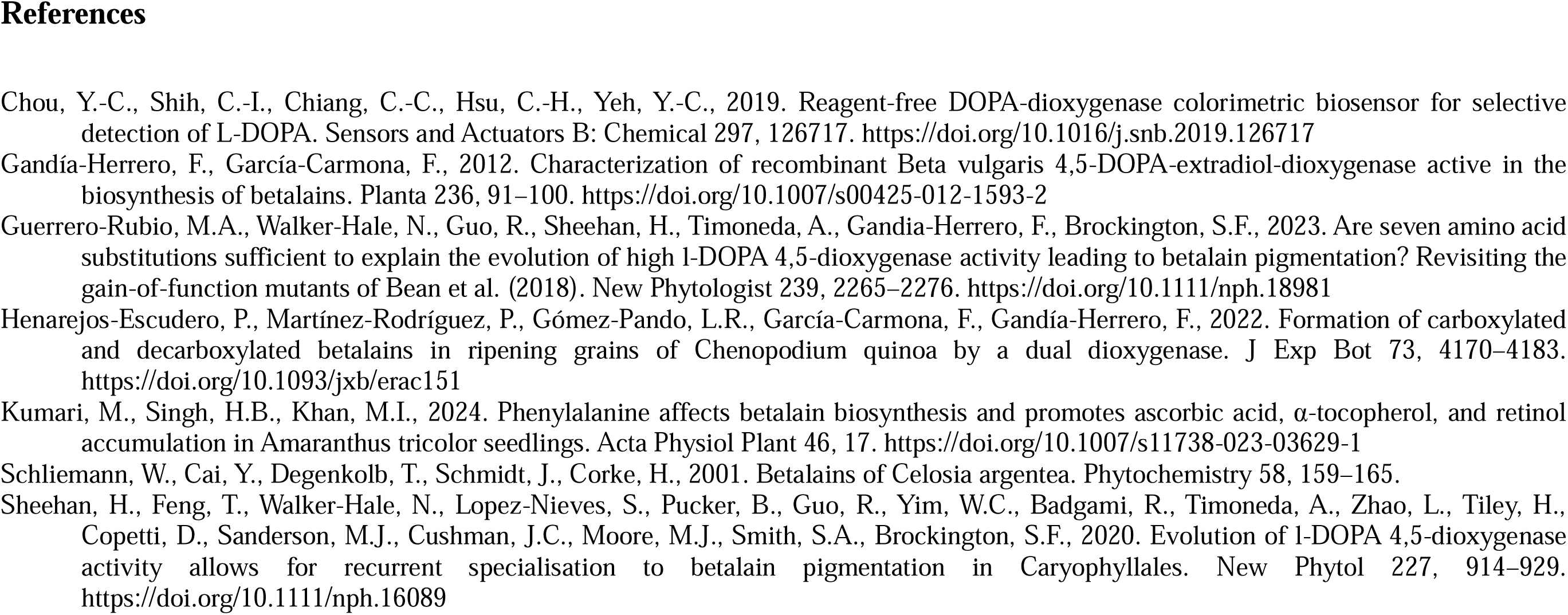
Grouping of LigB homologs of betalain-accumulating plants.

## 4. Discussion

Many LigB homologs from betalainic and non-betalainc plants have been reported (Chang et al., 2021; Chou et al., 2019; Christinet et al., 2004; Chung et al., 2015; Gandía-Herrero and García-Carmona, 2012; Guerrero-Rubio et al., 2023; Henarejos-Escudero et al., 2022; Li et al., 2024; Sasaki et al., 2009; Sheehan et al., 2020; Takahashi et al., 2015, 2009). The betalain-biosynthetic LigB homolog, *DOD*, from *Basella alba* L. var. ‘Rubra’ has not been characterized yet. In the present study, we undertook RACE-PCR-based gene mining and amplification of the full-length candidate gene, *BrDOD1*. From the deduced amino acid sequence of BrDOD1, the predicted molecular weight was ∼30 kDa, which is in accordance with the other reported DODs from betalainic plants (Chou et al., 2019; Gandía-Herrero and García-Carmona, 2012; Guerrero-Rubio et al., 2023; Henarejos-Escudero et al., 2022). After Ni-NTA-based affinity chromatography purification of the expressed protein in *E. coli* BL21-CodonPlus cells, a very low activity was detected during the enzyme assay of the recombinant BrDOD1. This may be due to the loss of metal ions and oxidation of Fe^2+^ during the extraction and processing of the enzyme. In enzyme assays of betalain biosynthetic plant DODs, it has been reported that adding Fe^2+^ in the reaction mixture enhances the BA formation (Chiang et al., 2025; Sasaki et al., 2009). However, in other studies, the activity was detected without Fe^2+^ addition as well (Gandía-Herrero and García-Carmona, 2012; Sasaki et al., 2009). During aerobic protein extraction and purification processes, the cofactor (Fe^2+^) present in the native protein may be leached, thereby reducing the enzyme activity (Baier et al., 2015; Barry and Taylor, 2013). In agreement with this, Colabroy et al. (Colabroy et al., 2019, 2008) observed that the protein purified from a heterologous host lacked any activity due to iron loss during purification, requiring a reconstitution step to activate it. So, the addition/preincubation of the enzyme with Fe^2+^ in the reaction mixture may be necessary to obtain an optimum reaction condition. On the flip side of it, Fe^2+^ may promote complex formation with the substrate (L-DOPA) triggering its oxidation (Bijlsma et al., 2020) resulting in the reduction of the net substrate concentration that could be catalyzed for BA formation, thereby hindering enzyme-specific activity. In addition, the L-DOPA degradation products may inhibit the enzyme activity (Colabroy et al., 2008). Therefore, care must be taken while adding Fe^2+^ to the reaction mixture. On the other hand, oxidation of the active site metal ion (Fe^2+^) due to oxygen binding to it before enzyme assay or during purification using aerobic buffers may hinder the assay. Therefore, to overcome such inhibition and substrate oxidation, we performed DOD enzyme re-activation before enzyme assay by reconstituting the enzyme with reducing agents, metal ions (Fe^2+^) and also by flushing N_2_ gas to minimize the exposure to atmospheric oxygen and incubating for 30 minutes at 4° C (Colabroy et al., 2019). We recommend that future kinetic characterization studies should take into consideration the enzyme activation step to streamline (or standardize) the assay protocol to generate truly comparative steady-state kinetic parameters.

From the steady-state kinetics, it was clear that BrDOD1 had almost comparable *K*_M_ toward substrates L-DOPA and dopamine, but around 6.6-fold higher catalytic efficiency (*k*_cat_/*K*_M_) for L-DOPA (9.9 ± 0.2 nM^-1^ min^-1^) than dopamine (1.5 ±0.03 nM^-1^ min^-1^) (Table 1B). BrDOD1 is the first enzyme among the DOD1 group (see the proposed classification in Table 3) whose *K*_M_ has been determined for both L-DOPA and dopamine. The use of dopamine as a substrate for high-activity DOD1 is justified because of its analogous structure to L-DOPA and available reports on the presence of dopamine and its derived pigments in betalain-containing plants (Kumari et al., 2024; Martínez-Rodríguez et al., 2024; Schliemann et al., 2001). Logically, this may imply that, in the absence of L-DOPA or higher dopamine content in the plant, DOD1 group enzymes may act on dopamine, help remove oxidative stress caused by dopamine accumulation (Kostyn et al., 2020), and play a physiological role in stress response. Among the DOD1 group, BrDOD1 seem to have comparable affinity for L-DOPA with MjDOD1 (Chou et al., 2019) but lower *K*_M_ than BvDODAα1 (Guerrero-Rubio et al., 2023) (Table S4). MjDOD1 has also been shown to accept dopamine to form decarboxylated betalain (Lin and Yeh, 2017), though kinetic parameters are yet to be determined. A DOD2 homolog (as per the classification in Table 3) from *C. quinoa* was reported to show higher affinity for dopamine (*K*_M_ 0.19 mM) than L-DOPA (*K*_M_ 1.5 mM) (Henarejos-Escudero et al., 2022) (Table S4). Therefore, we conducted MD simulation studies involving BrDOD1-L-DOPA and BrDOD1-dopamine complexes and established the preference for L-DOPA as the substrate of BrDOD1 based on the stable RMSD plot of BrDOD1-L-DOPA complex than the BrDOD1-dopamine complex (Fig. 3A and B). More such studies are needed to understand the substrate preference of DOD1 and DOD2 group proteins to strengthen the proposed classification of DOD homologs (Table 3).

Another important point in the DOD enzyme assays is the inclusion of 10 mM ascorbic acid (Chiang et al., 2025; Chou et al., 2019; Gandía-Herrero and García-Carmona, 2012; Guerrero-Rubio et al., 2023; Henarejos-Escudero et al., 2022) to maintain the reduced state of the metal ion (Fe^2+^) in the active site (Sasaki et al., 2009). To further understand the role of ascorbic acid in the steady-state kinetics of BrDOD1 during BA and 6-decarboxy-BA formation, we performed the reactions under the same conditions but in the absence of 10 mM ascorbic acid. We observed that in the absence of 10 mM ascorbic acid both substrates showed substrate inhibition with *K*_i_ of 7552.5±552.8 *µ*M for L-DOPA and 3493±367.5 *µ*M for dopamine Table 1b). However, those inhibitory substrate concentrations are very high and may not have any implications in the cellular environment of *B. alba* L. var. ‘Rubra’, as evident from the physiological concentrations of L-DOPA and dopamine in the mature/ripe fruit pulp (Table 2; Table S3). In contrast, substrate inhibition was not detected in the presence of ascorbic acid, but the *K*_M_ and *V*_max_ were increased. Among the betalain-biosynthetic DOD enzymes, BrDOD1 is the first to be assayed in the presence and absence of cosolute ascorbic acid, whose presence in high concentration relative to the substrate and the enzyme, raised the *K*_M_ and *V*_max_. Such changes are common in the case of protein crowders (Alfano et al., 2024; Silverstein, 2019). Such cosolutes have differential effects on the *V*_max_ above or below the transition point substrate concentration ([S]_=_) (Silverstein, 2019) (Fig. 2E and Fig. 2F; Fig. S6A and Fig.S6B). The effect was inhibitory below [S]_=_, while above this concentration, the reaction rate increased. Such effects may be ascribed to the molecular crowding effect of 10 mM ascorbic acid (Aumiller et al., 2014), in addition to its role to maintain the reduced state of Fe^2+^. The same phenomenon may be happening in the cellular environment as well because of the abundance of ascorbic acid in *B. alba* L. var. ‘Rubra’ (Table 2), thereby acting as a cosolute inducing molecular crowing effects on BrDOD1 and maintaining the reduced state of the metal ion (Fe^2+^). In view of this, catalytic activity assays of betalain-biosynthetic DODs should be done with optimized ascorbic acid concentration in future to obtain steady-state kinetic parameters free from the molecular crowding effect.

In the present study, three full-length LigB homologs viz. BrDOD1, BrDOD2, and BrLigB having different physicochemical properties have been isolated for the first time from *B. alba* L. var. ‘Rubra’. We have classified LigB homologs in betalainic plants into three groups: high BA-forming activity or DOD1 group, marginal/low BA-forming activity or DOD2 group, and LigB group having low/marginal BA-forming activity (Table 3). In a previous study, a classification of DOD homologs based on BA-forming activity was reported (Sheehan et al., 2020). In addition to the BA-forming activity, we have used phylogenetic tree, structural features, particularly the loop-forming region of the putative substrate-binding pocket near the active site, and pI (Fig. 4). Our phylogenetic tree is in agreement with previous findings suggesting low BA-forming activity DODAβ-clade as the ancestral proteins, which duplicated to evolve independently into high BA-forming proteins at least three times (Walker-Hale et al., 2025). Part of the evolution was in the loop-forming region, which contributed to the independent molecular convergence into high BA-forming activity (Walker-Hale et al., 2025) because it forms part of the substrate-binding pocket(Chiang et al., 2025). Therefore, in our classification, the loop-forming region is also one of the bases. Based on the sequence alignment of each group proteins, we propose updating of the conserved loop-forming consensus sequence of the substrate-binding pocket from H-P-(S/A)-(N/D)-x-T-P (Christinet et al., 2004) to H-P-[S/L/T]-D-[D/E]-T-P to account for all the enzymes having high BA-forming activity, and H-P-N-[N/S/G]-T-P for low/marginal activity enzymes. The corresponding consensus sequence of LigB group proteins is H-N-L-R, which is present in the sequences of LigB group proteins from betalainic plants as well (Sheehan et al., 2020), though earlier it was considered to be present in non-betalainic plants only (Christinet et al., 2004) (Table 3; Fig. S8B). Further, our proposed classification is supported by the median of pI values of LigB homologs as well (Fig. 4C). A correlation between the median pI in the present study could be established with assay pH in previous studies, i.e., optimum pH was low (pH < 7) for enzyme assays of DOD1 group (low median pI, in this study) proteins, such as MjDOD1 and BvDODAα1 (Chou et al., 2019; Guerrero-Rubio et al., 2023), compared to DOD2 group (higher median pI, in this study) proteins such as BvDODAα2 (Gandía-Herrero and García-Carmona, 2012). However, the assay pH of CqDOD of DOD2 group did not show the correlation (Henarejos-Escudero et al., 2022). Further, even within DOD2 group, the clade in which CqDOD falls is diverged from that of BvDODAα2 and others (Fig. 4B). Such observations cannot be explained clearly, as only two aforementioned DOD2 group proteins have been characterized so far. On the other hand, no LigB group protein has been characterized yet from betalainic plants. Considering the higher median pI, LigB group proteins’ activity may be assayed at a higher optimum pH than other DOD groups. Even though pI has no strong overall relation to protein evolution, through the indels, pI gets shifted in ortholog proteins (Alendé et al., 2011). The difference in the pI of DOD homolog groups may be due to the amino acid substitution during evolution. Proteomes in archaea, bacteria, and eukaryota have different net charge densities, with archaea proteins carrying negative charge, while bacterial proteins carry slightly positive charge (or basic in nature than eukaryote) (Vallina Estrada and Oliveberg, 2022). LigB group proteins with higher pI (toward the basic side) in our classification, could be indicating its bacterial origin or ancestral LigB copy in plants. On the other hand, DOD1 group (high BA-forming activity) has a lower pI (acidic) than the other two groups, probably due to evolutionary selection toward producing more efficient enzymes to utilize the pool of substrate metabolite (L-DOPA, which is acidic) in betalain-accumulating plants.

## 5. Conclusion

Betalain biosynthetic L-DOPA/dopamine-4,5-dioxygenase from *Basella alba* L. var. ‘Rubra’ (BrDOD1) catalyzes the aromatic ring-opening of L-DOPA and dopamine to form betalamic acid and 6-decarboxy-betalamic acid, respectively, with better preference for L-DOPA than dopamine. Activation with metal ion treatment post purification restores the activity of purified recombinant BrDOD1. Further, inclusion of ascorbic acid in the enzyme assay completely overcame the substrate inhibition. In addition, ascorbic acid produced a crowding effect possibly by inducing better conformational stability and promoting substrate-catalyst interaction near or within the substrate-binding pocket and active site of BrDOD1 resulting in increased *K*_M_ and *V*_max_ values. Further studies on the role and mechanism of ascorbic acid on BrDOD1 (or other DODs) kinetics are warranted. There are three LigB homologs in *B. alba* L. var. ‘Rubra’. All plant LigB homologs may be divided into DOD1, DOD2, and LigB groups based on sequence homology, phylogenetic resolution, structural features, betalamic acid-forming activity, and physicochemical properties. As a diagnostic structural feature of the three plant LigB homolog groups, we propose the characteristic conserved motif sequences H-P-[S/L/T]-D-[D/E]-T-P, H-P-N-[N/S/G]-T-P, and H-N-L-R at the loop forming region forming part of the substrate-binding pockets of DOD1, DOD2, and LigB groups, respectively.

## Supporting information

Manuscript

## Author CRediT Statement

HBS: Data curation, Investigation, writing − original draft, Writing − review and editing. MIK: Conceptualization, Data curation, Formal analysis, Funding acquisition, Investigation, Project administration, Resources, Supervision, writing − original draft, Writing − review and editing.

## Acknowledgements

HBS is grateful to the Department of Biotechnology, HRD-PMU, Govt. of India, for granting DBT-JRF fellowship (DBT/2022-23/GU/1890) and MIK is grateful to DBT (BT/PR16902/NER/95/422/2015 and BT/PR51321/NER/95/1994/2023) and SERB (ECR/2016/000952), Govt of India, respectively, for supporting this work. We acknowledge the contribution of Mr. Himangshu Sarma of the Biotechnology department, Gauhati University in MD simulation. We also acknowledge Ashok Verma of ATCREC, Mumbai and P. Giridhar of CFTRI, Mysuru for providing the *E. coli* strains. We also express our gratitude to the Head of the Department of Biotechnology, Gauhati University for his encouragement, and allowing access to analytical facilities in the department, including at the Institutional Biotech Hub (IBH), Gauhati University.

## Appendix A. Supplementary information and data

This article contains supporting information as supporting data. It will be available in the supplementary information section of this manuscript.

## Data availability

All the sequence data of *Basella alba* L. var. ‘Rubra’ LigB homologs were submitted to NCBI GenBank database and granted accession numbers (MK605278.1, PX763555, PX763556, PX763557, PX763558, and PX763559). And LigB homologs used for phylogenetic tree construction and analysis of pI distribution are being provided as additional supporting data file S1 and S2.

