## Supplementary material for "*Basella alba* L. var. ‘Rubra’ L-DOPA/dopamine-4,5-dioxygenase 1 prefers L-DOPA over dopamine and ascorbic acid enhances its activity": Manuscript

**1. Supplementary methods**

**1.1. Spectrophotometric analysis of betalains**

Matured pulp of *B. alba* L. var. ‘Rubra’ was collected and after the removal of seed, the pulp portion was ground in acidified water (pH 5.5) and supernatant was collected by centrifugation at 8000 ×*g* for 10 min. The pellet was again re-extracted with acidified water and supernatant was collected by centrifugation and analyzed using a spectrophotometer (UV-1800, M/s Shimadzu Corp., Kyoto, Japan).

**1.2. HPLC analysis and quantification of metabolites**

Mature fruits of *B. alba* L. var. ‘Rubra’ were collected and after the removal of seeds manually, pulps were ground in the 10 mL of extraction medium consisting of 0.05% TFA; 5% acetonitrile in H_2_O (v/v) (pH adjusted to 2.0 with phosphoric acid) and the supernatant was collected after centrifugation for 8000 ×*g* for 10 min. Again, the pellet was reextracted with 10 mL of the same extracted medium and supernatant was pooled together. Then, the analysis was done using a slight modification of a reported HPLC protocol (Zhou et al., 2012). An aliquot of 20 *μ*L of the sample was injected into the HPLC (Shimadzu LC20AD) equipped with a photo-diode array (PDA) detector and separated in a YMC reversed phase column (250×4.6 mm, 5 *μ*m particle size and 12 nm pore size). The reaction mixture was separated at a 1.2 mL/min flow rate with an isocratic condition for 15 min. And the quantification was done using the respective standard calibration curves of L-DOPA and dopamine.

For ascorbic acid analysis, sample was ground in 10 mL of extraction solvent (0.05% TFA; 5% acetonitrile in H_2_O (v/v) pH adjusted to 2 with phosphoric acid) and again re-extracted with 5 mL of extraction medium and the supernatant was pooled together and 20 *μ*L of sample were injected into the HPLC (Shimadzu LC20AD) equipped with a photo-diode array (PDA) detector and separated in a YMC reversed phase column (250×4.6 mm, 5 *μ*m particle size and 12 nm pore size). The reaction mixture was analyzed at a 1.2 mL/min flow rate with an isocratic condition for 15 min. And the quantification was done using a standard calibration curve ascorbic acid.

**1.3. Structure modelling, docking and MD simulation**

3D-structure of BrDOD1 protein was modelled using the deduced amino acid sequence on the Alphafold3 server (https://alphafoldserver.com/) and the best-fit model was selected and validated using the PROCHECK tool (Laskowski et al., 1993) of SAVESv6.1 server (https://saves.mbi.ucla.edu/) through Ramachandran plot analysis. After validation, molecular docking was performed on CB-Dock2 (Liu et al., 2022) server (https://cadd.labshare.cn/cb-dock2/php/index.php) and the protein-ligand complex scoring the lowest energy was selected, visualized, and interacting residues were identified in Discovery Studio 2021. Molecular dynamics (MD) simulations were performed using the Desmond module v7.6 (Academic License; D. E. Shaw Research) integrated within the Schrödinger Maestro v 13.8 interface (Schrödinger Release 2023-24). Desmond is widely used for biomolecular simulations due to its high-performance parallel algorithms and validated force-field implementation. Each protein–ligand complex was prepared using the System Builder on Maestro. The systems were solvated with the TIP3P explicit water model inside an orthorhombic box, maintaining a 10 Å buffer around the solute. Charge neutrality was ensured by adding Na⁺ or Cl⁻ counterions, followed by the addition of 0.15 M NaCl to mimic physiological ionic strength. Parameterization of both protein and ligand was carried out using the OPLS-2005 force-field. Short-range non-bonded interactions were evaluated with a 9 Å cutoff, while long-range electrostatics were computed using the particle mesh Ewald (PME) method. All simulations were conducted under NPT ensemble conditions at 298 K and 1.01325 bar. Temperature coupling was maintained using the Nosé–Hoover thermostat, and pressure was controlled using the Martyna–Tobias–Klein barostat. Before production runs, each solvated system underwent 200 ps of energy minimization and relaxation following the default Desmond equilibration protocol. The final production MD trajectory was recorded for 100 ns for each complex with the integration time step at 2 fs to evaluate structural stability and ligand-binding dynamics.

**1.4. In-silico analysis of LigB homologs**

After sequencing and confirmation, the nucleotide sequence of full-length putative LigB homologs of *B. alba* L. var. ‘Rubra’ were used to deduce their corresponding protein sequences via the NCBI ORFfinder tool using the confirmed nucleotide sequences (https://www.ncbi.nlm.nih.gov/orffinder/). These amino acid sequences were then aligned using MUSCLE5 (Edgar, 2022), and the secondary structure alignment was conducted on the ESPript server (https://espript.ibcp.fr/ESPript/cgi-bin/ESPript.cgi) (Robert and Gouet, 2014) using the aligned LigB homolog sequences as input against the 3D structure of *Beta vulgaris* DODA*α*2 (PDB ID: 8IN2) (Chiang et al., 2025). Further, the deduced amino acid sequences were used to predict the theoretical isoelectric point (pI), molecular weight (Kozlowski, 2016), and subcellular localizations using the DeepLoc 2.0 web server (Thumuluri et al., 2022).

**1.5. Construction of phylogenetic tree of LigB homologs**

Amino acid sequence homologs of LigB were searched from plants within the order Caryophyllales against the NCBI database (https://blast.ncbi.nlm.nih.gov/Blast.cgi) using the BLASTp program. The BLAST output sequences with a query coverage ranging from 75% to 100% and an identity of at least 50%, were retrieved. Redundant sequences were removed using the Jalview tool (Procter et al., 2021). Subsequently, the sequences 273 Caryophyllales LigB homologs (Supplementary data S1) were aligned using MUSCLE5 (Edgar, 2022), and an unrooted phylogenetic tree was constructed using the best-fit model Q. plant+R7 according to Bayesian Information Criterion (BIC), with 100,000 ultrafast bootstrap replicates in the IQTREE2 tool (Nguyen et al., 2015).

**1.6. Identification of conserved motifs in LigB homologs and their distribution**

Further, for conserved motif analysis, from the 273 Caryophyllales “LigB homologs” sequences were visually inspected for the presence or absence of specific conserved amino acid sequence motifs (Chiang et al., 2025; Christinet et al., 2004; Li et al., 2024) in the aligned data to identify group-specific conserved motifs associated with reported betalamic acid-forming activities. Additionally, percent abundances and distribution across the betalainic plants of Caryophyllales were determined for each LigB homolog group having a specific conserved motif region.

**1.7. Computation of isoelectric point of LigB homologs**

After grouping and identifying conserved motifs of LigB homologs, the theoretical isoelectric point (pI) of different LigB homolog protein sequences viz. DOD1 (58), DOD2 (97), and LigB (72) proteins (Supplementary data S2) was computed using the Isoelectric Point Calculator (IPC) (Kozlowski, 2016) web server (http://isoelectric.org/) and the average isoelectric points were plotted in a box plot graph using Origin 8.5.

**Supplementary tables and legends**

**Table S1A.** Primers used in the amplification of partial LigB homologs

| Sl no | Primer name | Sequences (5′--------- 3′) |
| --- | --- | --- |
| 1 | Partial DOD1-like Forward | GTTCTACCTGTCGCATGG |
| 2 | Partial DOD1-like Reverse | CTTCATCTGGAACATGCAAG |
| 3 | Partial DOD2-like Forward | GATGAGGATCGAGGCAAGG |
| 4 | Partial DOD2-like Reverse | GGCCAACATCCCTGTATGTC |
| 5 | Partial LigB-like Forward | ATCGATTCTCATCATCTCCGG |
| 6 | Partial LigB-like Reverse | GTATCCTGAGGCCAACATCC |

**Table S1B.** Primers used in RACE-PCR for the amplification of DOD1 3′ and 5′ cDNA ends

| Sl. No. | Primer name | Sequences (5′--------- 3′) |
| --- | --- | --- |
| 1 | 5̕̕ DOD1 GSP1 | CTTCATCTGGAACATGCAAG |
| 2 | 5̕̕ DOD1 GSP2 | TGGCCAGCAGATACAGAG |
| 3 | 5̕̕ DOD1 GSP3 | ACTTGGGTTTGATGGGGAAC |
| 4 | 3̕ DOD1 GSP1 | GTTCTACCTGTCGCATGG |
| 5 | 3̕ DOD1 GSP2 | CATGGGAATCCGGCAATG |
| 6 | Anchored primer | GACCACGCGTATCGATGTCGACTTTTTTTTTTTTTTTTV |
| 7 | Oligo dT anchored | GACCACGCGTATCGATGTCGAC |

**Table S1C.** Primers used in the full-length amplification of LigB homologs

| **Sl. No.** | **Primer name** | **Sequences (5′--------- 3′)** |
| --- | --- | --- |
| 1 | DOD1 Forward | GGTGATGATGATGACAAGATGGGTGTTGGGAAGCAAATG |
| 2 | DOD1 Reverse | GGAGATGGGAAGTCATTATGGAGTCAAGTGGAAGTGAAC |
| 3 | DOD2 Forward | GGTGATGATGATGACAAGATGGCCGACCTCTCACAG |
| 4 | DOD2 Reverse | GGAGATGGGAAGTCATTAGCAGGAAGTGAACTTGTAAGAGG |
| 5 | LigB Forward | GGTGATGATGATGACAAGATGACGAGGAAAAAACCCAAGTTGA |
| 6 | LigB Reverse | GGAGATGGGAAGTCATTAATCAGCAGAAGTGAACTTATATGACGC |

**Table S2.** Substrate affinity and substrate affinity (*K*_M_) and catalytic efficiency of BrDOD1 in presence of ascorbic acid analyzed by Hanes-Woolf plot

| Substrate | *V*_max_ (*µ*M min^-1^) | *K*_M_ *(µ*M) | *k*_cat_ (min^-1^) | *k*_cat_/*K*_M_ (nM^-1^ min^-1^) |
| --- | --- | --- | --- | --- |
| L-DOPA | 5.5±0.16 | 202±10.6 | 1.9±0.05 | 9.3±0.24 |
| Dopamine | 1.6±0.04 | 372 ±13.1 | 0.54±0.02 | 1.5±0.02 |

Mean ± SD (n=4)

**Table S3**. L-DOPA and dopamine contents of betalainic plants on fresh/dry weight basis

| Sample | L-DOPA (*μ*g/g fw) | DPH (*μ*g/g fw) | Ascorbic acid (*μ*g/g fw) | Reference |
| --- | --- | --- | --- | --- |
| *Basella alba* L. var. ‘Rubra’ fruit (mature/ripe pulp) | 187.3 ± 5.2 | 71.1 ± 5.6 | 597.1 ± 18.9 | This study |
| *Amaranthus tricolor* red seedlings (12-day-old) | 2.2 ± 0.1 | 0.26 ± 0.03 | 21.6 ± 1.2 | (Kumari et al., 2023) |
| *Amaranthus tricolor* red seedlings (16-day-old) | 0.89 ± 0.02 | 0.32 ± 0.02 | 14.9 ± 1.0 |  |
| *Amaranthus tricolor* red seedlings (one month old) | 0.27 ± 0.04 | 0.14 ± 0.01 | 50.2 ± 2.3 |  |
| *Celosia cristata* (violet) callus culture | Not studied | 1.2 ± 0.07 | Not studied | (Lystvan et al., 2018) |
| *Celosia cristata* inflorescence (violet) | Not studied | 5.8 ± 2.1 ^#^ | Not studied |  |
| *Celosia plumosa* inflorescence (red) | Not studied | 5.8 ± 1.3 ^#^ | Not studied |  |
| *Celosia plumosa* inflorescence (orange) | Not studied | 5.1 ± 1.7 ^#^ | Not studied |  |
| *Celosia argentea* var. cristata inflorescence (yellow) | Not studied | 6.3 * | Not studied | (Schliemann et al., 2001) |
| *Hylocereus polyrhizus* red flesh fruit | L-DOPA >> DPH | | Not studied | (Suh et al., 2014) |
| *Hylocereus undatus* white flesh fruit | L-DOPA >> DPH | | Not studied |  |

* mg/g fresh weight; DPH-dopamine

^#^ mg/g dry weight

**Table S4.** Reported substrate affinity (*K*_M_) and catalytic efficiency of functional plant DODs

| Candidate DOD | *K*_M_  for L-DOPA (mM) | *K*_M_  for Dopamine (mM) | L-DOPA *k*_cat_/*K*_M_  (*μ*M^-1^min^-1^) | Dopamine  *k*_cat_/*K*_M_  (*μ*M^-1^min^-1^) | L-DOPA *K*_i_  (mM) | Dopamine *K*_i_  (mM) | Reference |
| --- | --- | --- | --- | --- | --- | --- | --- |
| BvDODA*α*1 | 2.73 ± 9.558 | n.a. | n.a. | n.a. | 0.0281 ± 0.09 | n.a. | (Guerrero-Rubio et al., 2023) |
| MjDOD1 | 0.168 | 1.09 | 7.54×10^-3^ | n.a. | n.a. | n.a. | (Chou et al., 2019) |
| BvDODA*α*2 | 6.9 | n.a. | 1.38×10^-5^ | n.a. | n.a. | n.a. | (Gandía-Herrero and García-Carmona, 2012) |
| CqDOD2 | 1.5 ± 0.01 | 0.19 ± 0.05 | n.a. | n.a. | n.a. | n.a. | (Henarejos-Escudero et al., 2022) |

n.a.- not available.

**Table S5.** In-silico analysis of LigB homologs of *Basella alba* L. var. ‘Rubra’

| DOD/LigBs | Length (bp) | Length (aa) | Isoelectric point (pI) | Mol. wt. (kDa) | Subcellular localization |
| --- | --- | --- | --- | --- | --- |
| BrDOD1 | 813 | 270 | 4.99 | 30.214 | Cytoplasm and Nucleus |
| BrDOD2 | 792 | 263 | 5.54 | 29.011 | Cytoplasm |
| BrLigB | 996 | 331 | 8.59 | 37.645 | Endoplasmic reticulum |

**Supplementary figures and legends**

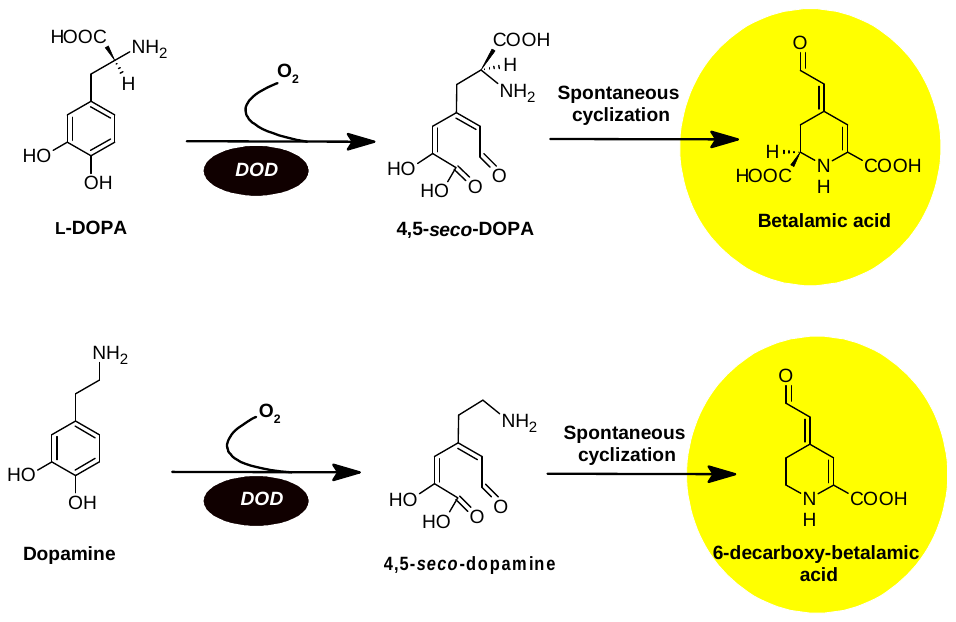

**Fig. S1**. Reaction scheme of betalamic acid and 6-decarboxy betalamic acid formation by DOPA-4,5-extradiol ring-cleaving dioxygenase (DOD1) involved in the betalain biosynthesis of plants (Adapted from Khan & Giridhar, 2015; Guerrero-Rubio et al., 2020).

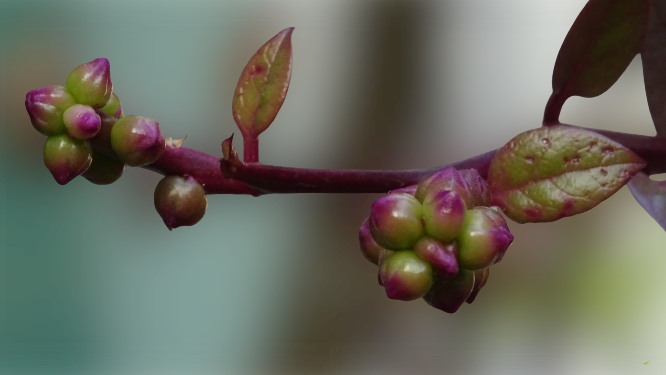

**Fig. S2**. A representative picture of the *Basella alba* L. var. ‘Rubra’ aerial part.

**(A)**

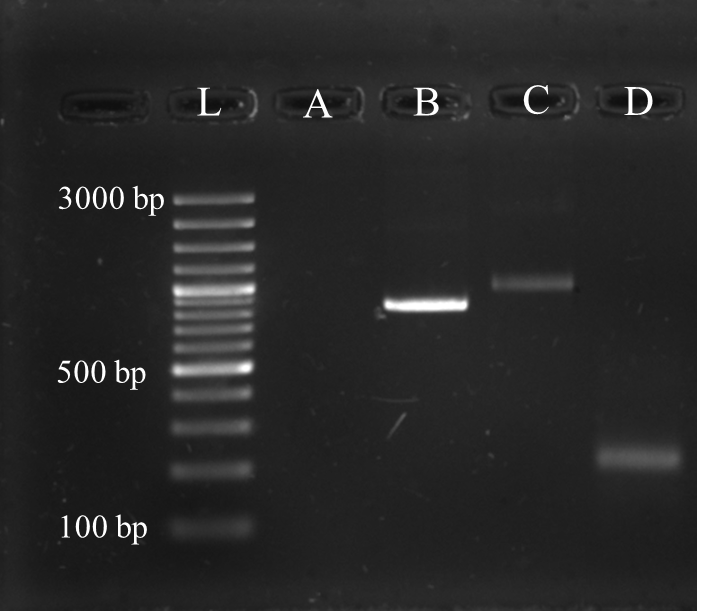

**(B)**

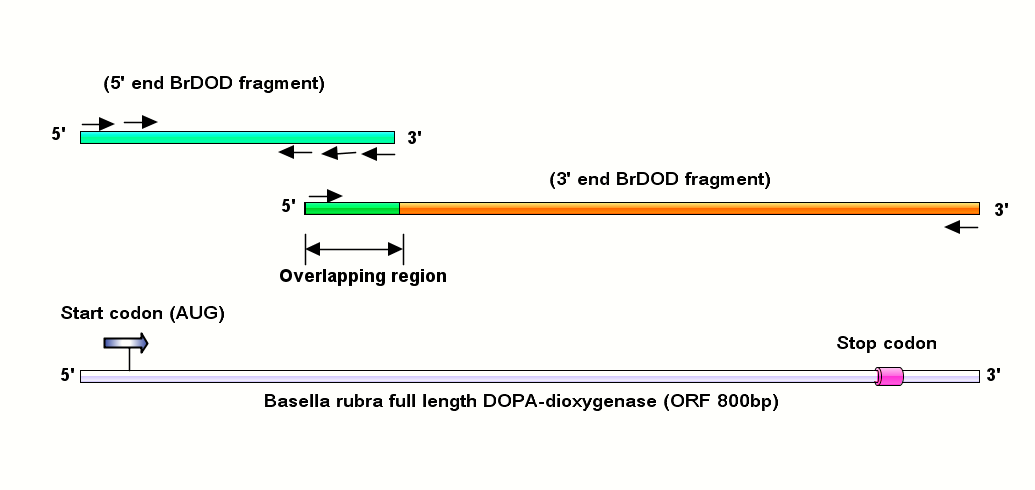

**Fig. S3.** RACE-PCR-cloned full-length LigB homolog BrDOD1. **A)** Gel picture showing the DNA band (L-100 bp ladder; A- blank; B- full-length BrDOD1; C- 3′ end BrDOD1; and D- 5′ end BrDOD1), and **B)** schematic representation of assembly of partial fragments into full-length BrDOD1,

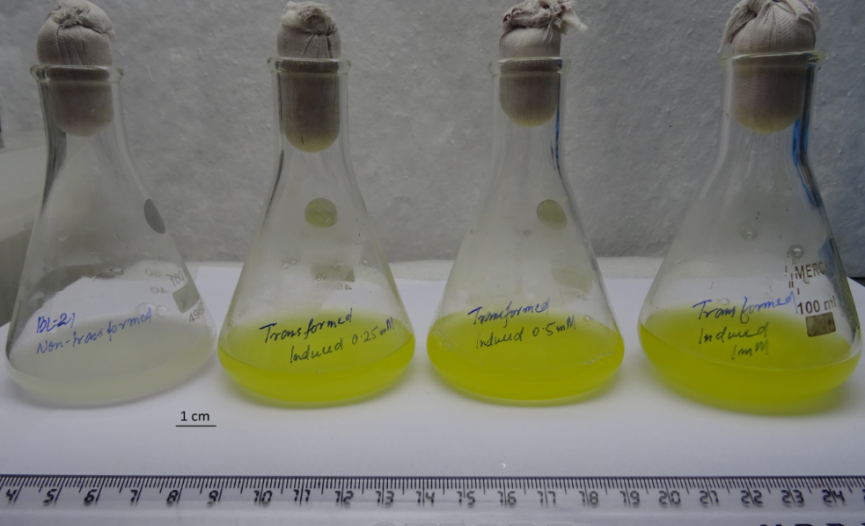

**Fig. S4.** Formation of betalamic acid by transformed *E. coli* BL21-CodonPlus cells expressing BrDOD1 after addition of L-DOPA.

**(A)	(B)**

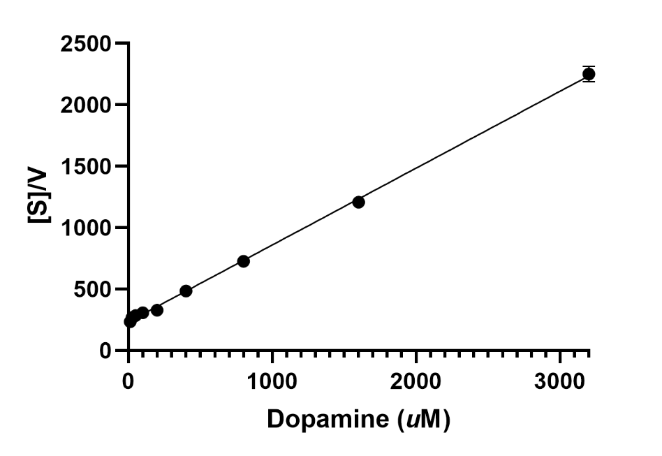

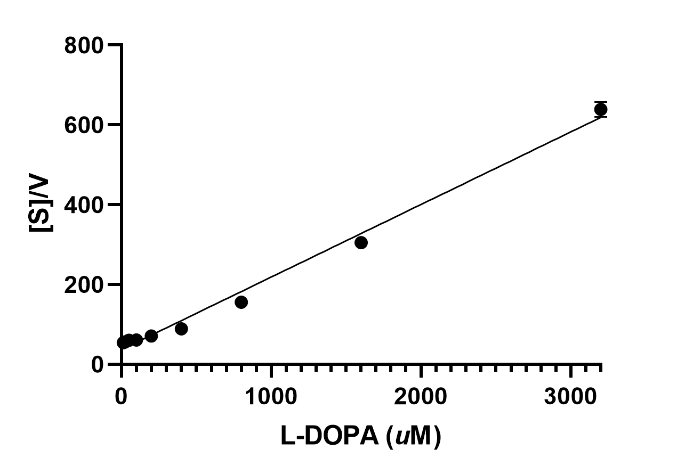

**Fig. S5.** Linear steady-state kinetic data of BrDOD1 fitted with Hanes-Woolf plot for different substrates. **A)** L-DOPA and **B)** dopamine.

**(A)	(B)**

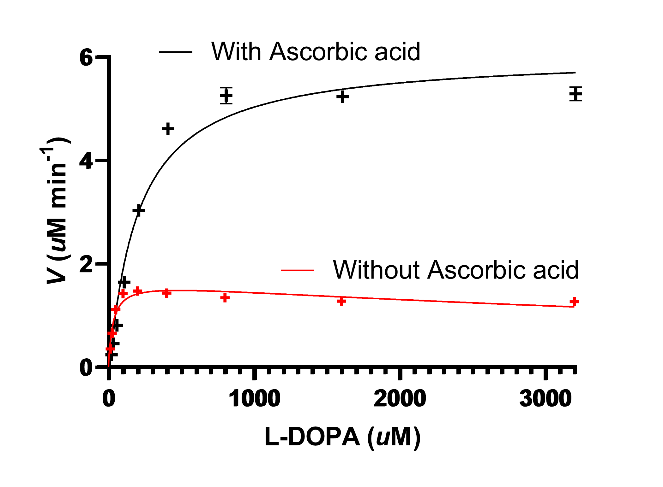

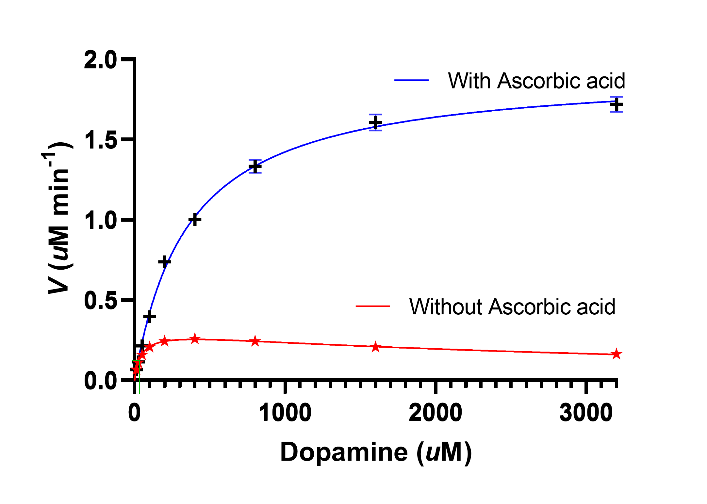

**Fig. S6.** Overlay of steady-state kinetic curves of BrDOD1 toward **(A)** L-DOPA and **(B)** dopamine in the presence or absence of ascorbic acid.

**(A)**

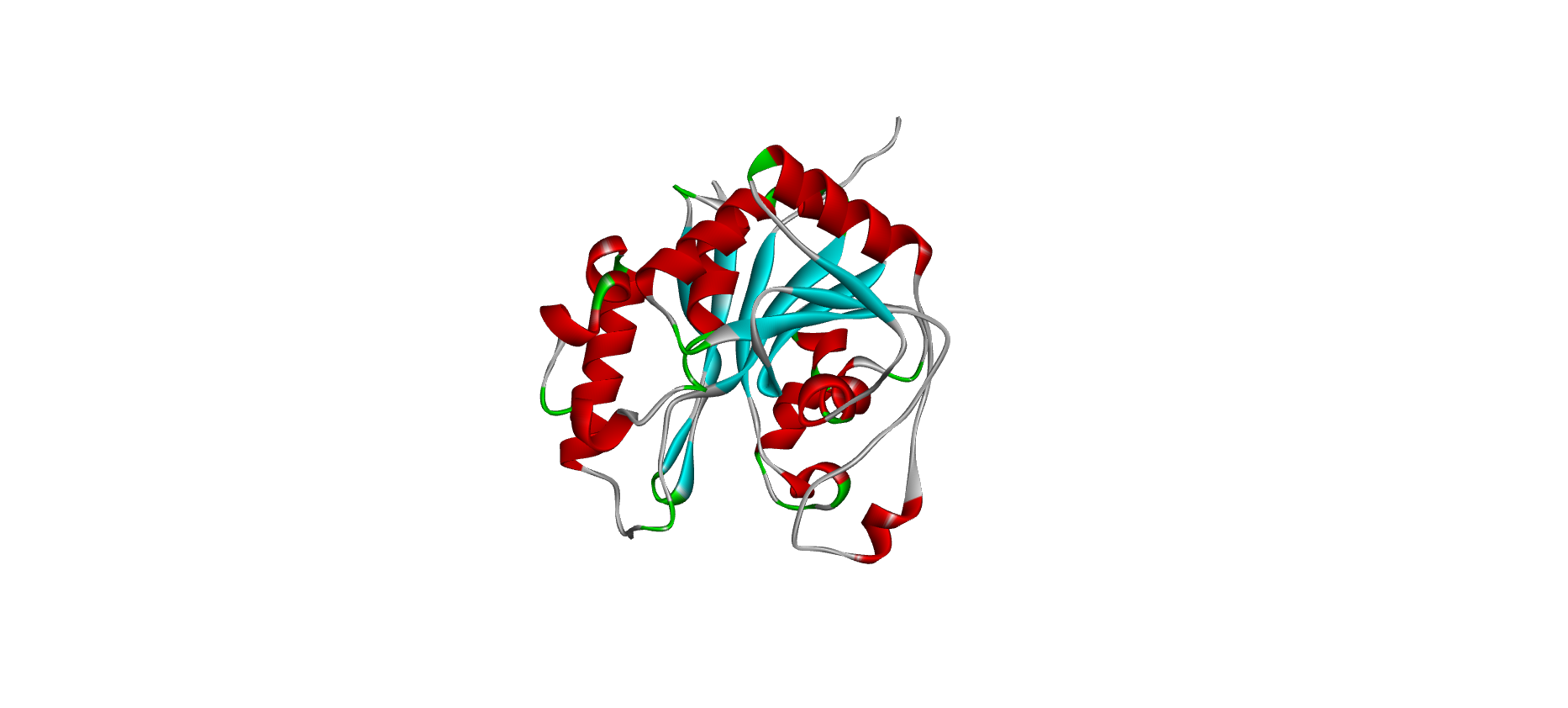

**(B)**

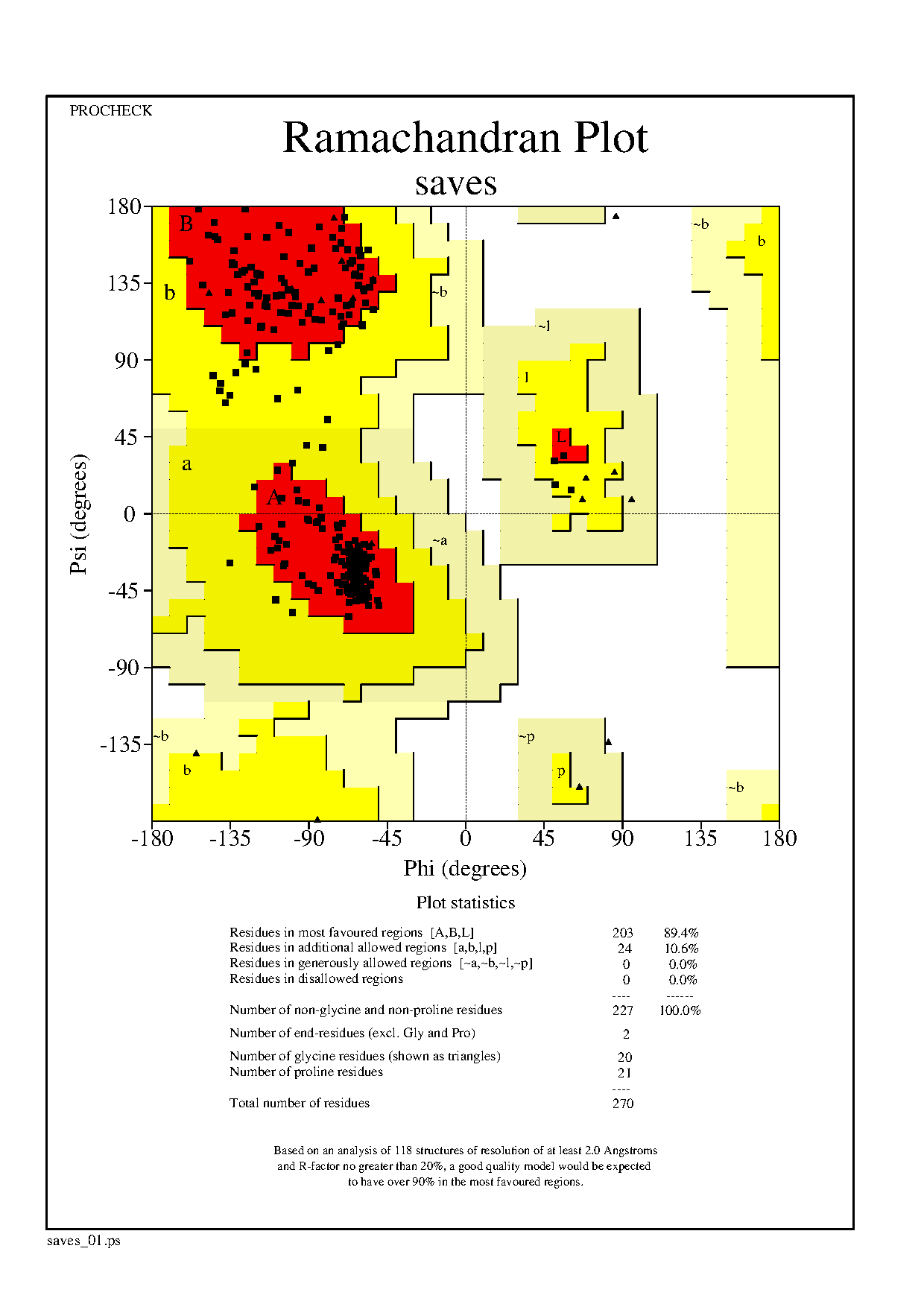

**(C)	(D)**

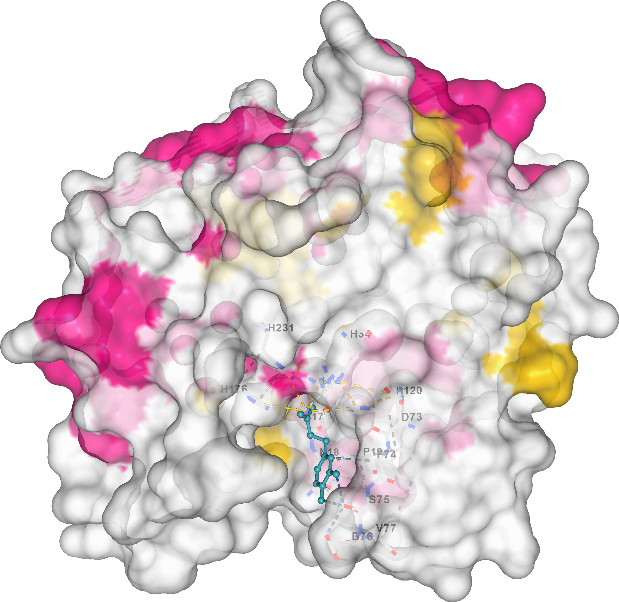

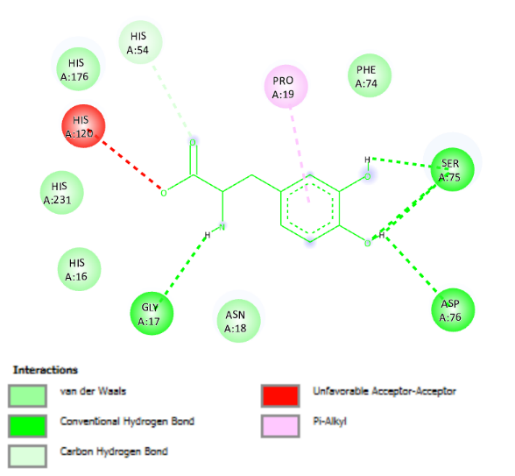

**(E)	(F)**

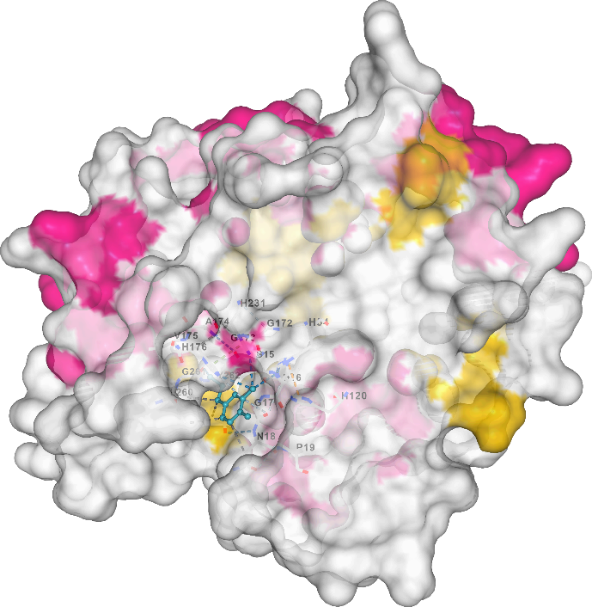

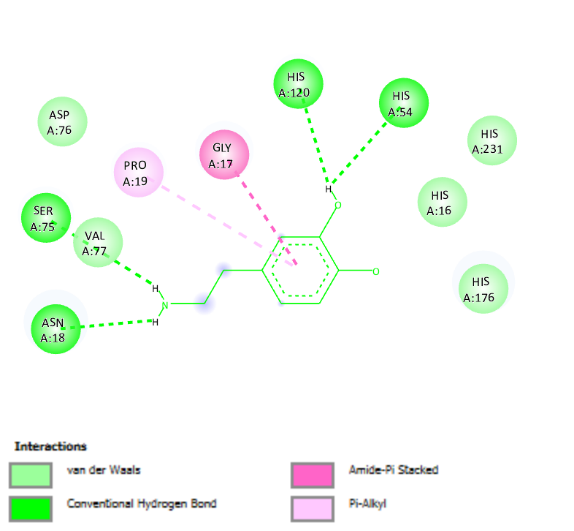

**Fig. S7.** Modelling of BrDOD1 and docking of L-DOPA and dopamine. **A)** AlphaFold3-generated 3-D model of BrDOD1, **B)** Ramachandran plot of BrDOD1 model, **C)** docked complex of BrDOD1 with L-DOPA, **D)** a section of the residues interacting with L-DOPA, **E)** docked complex of BrDOD1with dopamine, and **F)** a section of the residues interacting with dopamine.

**(A)**

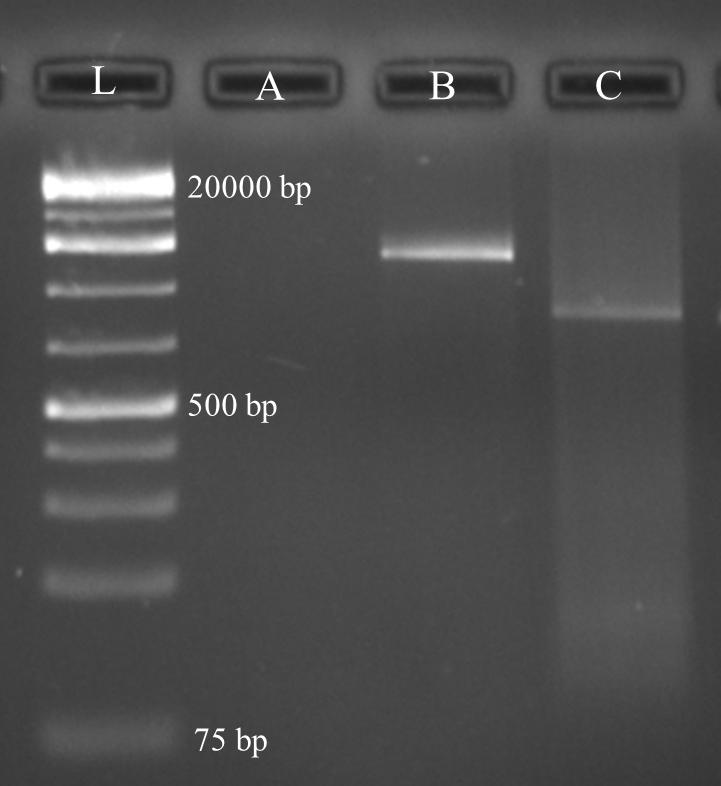

**(B)**

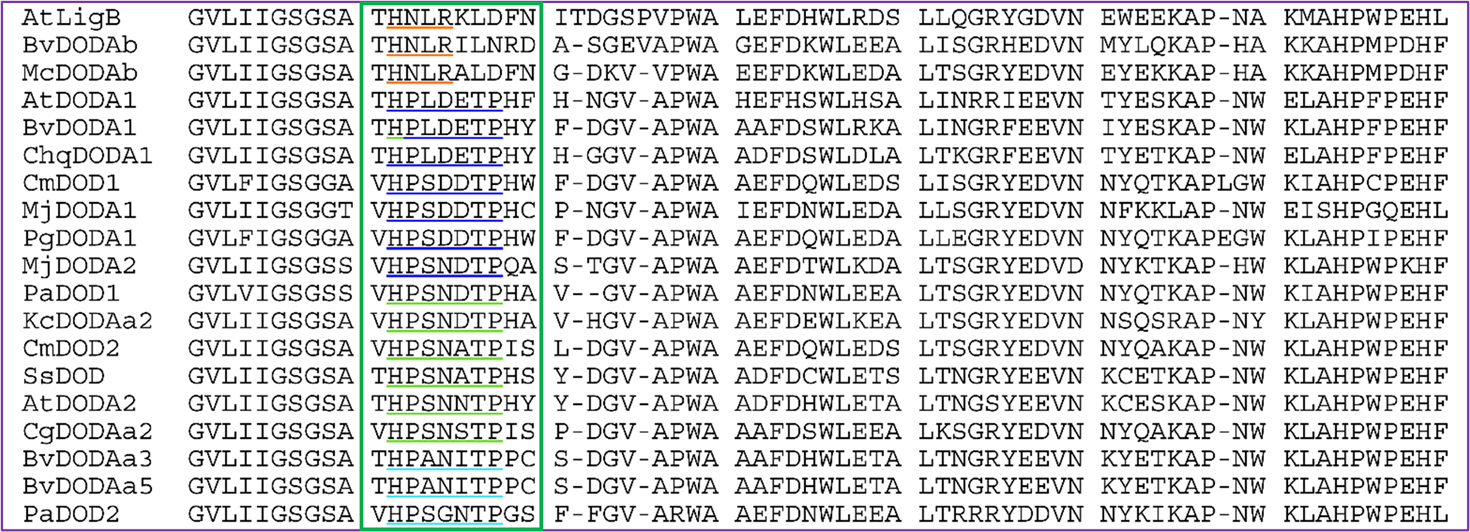

**Fig S8.** Second and third LigB homologs from *B. alba* L. var. ‘Rubra’ and sequence alignment of LigB homologs from different plants. **A)** Gel picture showing amplified full-length BrDOD2 and BrLigB PCR product (L- 1 kb plus ladder; A- bank; B- full-length BrDOD2; and C- full-length BrLigB) and **B**) a portion of sequence alignment using representative DOD protein sequences (accession IDs are given in Supplementary data S2) showing specific conserved motifs of each high BA-forming activity, low/marginal BA-forming activity DOD and non-BA-forming DOD groups.
